# Intensive Single Cell Analysis Reveals Immune Cell Diversity among Healthy Individuals

**DOI:** 10.1101/2021.10.18.464926

**Authors:** Yukie Kashima, Keiya Kaneko, Patrick Reteng, Nina Yoshitake, Lucky Ronald Runtuwene, Satoi Nagasawa, Masaya Onishi, Masahide Seki, Ayako Suzuki, Sumio Sugano, Mamiko Sakata-Yanagimoto, Yumiko Imai, Kaori Nakayama-Hosoya, Ai Kawana-Tachikawa, Taketoshi Mizutani, Yutaka Suzuki

**Affiliations:** Department of Computational Biology and Medical Sciences, Graduate School of Frontier Sciences, The University of Tokyo, Kashiwa, Chiba, Japan; Division of Collaboration and Education, International Institute for Zoonosis Control, Hokkaido University, Sapporo, Hokkaido, Japan; AIDS Research Center, National Institute of Infectious Disease, Tokyo, Japan.; Division of Breast and Endocrine Surgery, Department of Surgery, St. Marianna University School of Medicine, Kawasaki, Kanagawa, Japan; Institute of Kashiwa-no-ha Omics Gate, Kashiwa, Chiba, Japan; Future Medicine Education and Research Organization at Chiba University Inohana 1-8-1 Chuo, Chiba-city, Chiba, 260-8670 Japan; Department of Hematology, Faculty of Medicine, University of Tsukuba, Tsukuba, Ibaraki, Japan; Laboratory of Regulation for Intractable Infectious Diseases, National Institutes of Biomedical Innovation, Health and Nutrition (NIBIOHN), Osaka 567-0085, Japan

**Author notes:** These authors equally contributed (Yukie Kashima, Keiya Kaneko, Patrick Reteng). **Contact information** Correspondence should be addressed to: Yutaka Suzuki Laboratory of Functional Genomics, Department of Computational Biology and Medical Sciences, Graduate School of Frontier Sciences, The University of Tokyo, 5-1-5, Kashiwanoha, Kashiwa, Chiba 277-8562, JAPAN.

## Abstract

It is believed that immune responses are different between individuals and at different times. In addition, personal health histories and unique environmental conditions should collectively determine the present state of immune cells. However, the cellular and molecular system mechanisms underlying such heterogeneity remain largely elusive. In this study, we conducted a systematic time-lapse single-cell analysis, using 171 single-cell libraries and 30 mass cytometry datasets intensively for seven healthy individuals. We found substantial diversity in immune cell populations and their gene expression patterns between different individuals. These patterns showed daily fluctuations even within the same individual spending a usual life. Similar diversities were also observed for the T cell receptor and B cell receptor repertoires. Detailed immune cell profiles at healthy statuses should give an essential background information to understand their immune responses, when the individual is exposed to various environmental conditions. To demonstrate this idea, we conducted the similar analysis for the same individuals on the vaccination of Influenza and SARS-CoV-2, since the date and the dose of the antigens are well-defined in these cases. In fact, we found that the distinct responses to vaccines between individuals, althougth key responses are common. Single cell immune cell profile data should make fundamental data resource to understand variable immune responses, which are unique to each individual.

## INTRODUCTION

The human immune system consists of ingenious immune cells. It is widely known that the immune cells are collectively responsible for the versatile immune responses of an individual by shaping the immune landscape. The immune landscape should differ between individuals with distinct medical history and lifestyles, depending on genetic backgrounds and geographic origins. However, the influences and consequences of how immune cells maintain the previous memory of immune responses in a healthy state and respond to stimulation in normal life are largely unknown. It is partly because current knowledge on immune responses has been accumulated from laboratory animal models or diseased individuals. Even among individuals with a healthy appearance, infections, which may be mostly asymptomatic, occur daily.

Diverse immune responses, whether mild or severe, primarily occur at the infection site. However, it is commonly accepted that the immune profile is represented, at least in part, by circulating white blood cells i.e., peripheral blood mononuclear cells (PBMCs). Among the various cell types in PBMCs, innate immune cells are the first basal responders to antigen exposure, followed by the adaptive immune system. More precisely, monocytes are the first responder to various immune stimulations, the classical monocytes being CD14^+^ monocyte and the non-classical monotypes being CD16^+^ monocytes. When monocytes are recruited to the site of immune responses, they mature into macrophages or dendritic cells (DCs). DCs engulf antigens through phagocytosis and migrate to lymph nodes where they present the antigens to T cells, and an adaptive immune response is invoked. Natural killer (NK) cells are also recruited to secret proteins that kill the infected cells and trigger the adaptive immune response (Nicholson 2016).

The adaptive immune responses depend on the specific recognition of an antigen by T cells or B cells at its recognized part called epitope(Minervina et al. 2019). In T cells, the VDJ segments of the T cell receptor (TCR), consisting of alpha and beta chains, are imperative to function properly. A set of VDJ segments unique to each cell determines the sequence of the antigen-binding site presented on the cell surface. Cells sharing the same VDJ sequence are said to have the same “clonotype.” Effector CD8^+^ T cells, also known as cytotoxic CD8^+^ T cells, are activated upon antigen exposure via class I MHC molecule and leads the target cells to death (Golubovskaya and Wu 2016). Similarly, B cell receptors (BCRs) are composed of immunoglobulin molecules presented on the outer cell membrane. In B cells, antigen specificity is determined by the heavy and light chains of the immunoglobulins (IgK and IgL). Once exposed to an antigen, naïve B cells differentiate into either memory B cells or plasmablasts, which differentiate into plasma cells to produce antibodies.

Conventionally, flow cytometry and fluorescence-activated cell sorting, or more recently bulk RNA-seq, have been the standard methods for monitoring the states of PBMCs and the immune systems they represent(Stubbington et al. 2017). However, substantial concerns have been raised about these methods. The main drawback is that, although these methods offer high-throughput and extensive gene detection, the obtained data would come from the cell mass in bulk; hence, the gene expression for a particular cell population is not represented separately. Also, it has been impossible to analyze the VDJ patterns, which are unique to individual cells, especially in association with the status of their expressing immune cells. Single-cell RNA-seq (scRNA-seq), a recently introduced method, enables a detailed observation at the single-cell level(Stubbington et al. 2017). With this single-cell analytical approach, cellular heterogeneity previously masked using bulk RNA-seq is now open to investigation to assess the response of a certain cell to an identified antigen.

Using scRNA-seq, recent studies have illustrated in great detail how the immune cells practically change their profiles in response to disease and infection. For example, immune responses to infection with Severe Acute Respiratory Syndrome Coronavirus (SARS-CoV)-2, which is a pressing global health issue, has been analyzed very fervently last year. A study on the PBMCs of patients with moderate to severe coronavirus 2019 (COVID-19) reported that the relative abundance of naïve and activated T cells, mucosal-associated invariant T cells (MAIT), and monocyte-derived DCs decreased with disease severity, while T cells, plasma B cells, classical monocytes, and platelets increased (Zhang et al. 2020). Particularly, the timing and degree of induction of a subclass of T cells, called gamma delta T cells, appeared to be an important factor in determining the severity of the infections. Furthermore, it is noteworthy that all the cell populations except for the activated T cells were restored at convalescence.

In contrast to disease states, the extent to which immunity may differ amongst healthy individuals remains almost totally elusive, although fundamental information on which various immune responses occur should be available. Only a handful of studies have been reported. TCR-VDJ gene-targeting PCR analysis revealed that TCR repertoires of memory T cells are at least partially specific to individuals. In contrast, TCRs from naïve T cells showed no such individuality. The diverse immunological profiles, depending on individuals, may reflect their genetic background, infection history, and interactions with environments over a lifetime. However, how such diversity is acquired, maintained, and serves as a ground state for immune responses in generally healthy individuals remains almost totally unknown.

In this study, we describe our observations of the immune profile heterogeneity amongst healthy adults at the single-cell level, measured using scRNA-seq of the cellular transcriptomes and the VDJ repertoires. The profiles were further perturbed by vaccination against influenza and SARS-CoV-2. The resulting observations should be explained by the distinct “personal immunological landscape” shaped by each individual throughout their life and their present infection state, although participants reported a good health state during the study period. The broad aim of this study is to help understand baseline diversity in control groups regularly used in immunological studies of disease.

## RESULTS

### Generation and evaluation of the scRNA-seq data for healthy individuals

To characterize the immunological landscape of seven healthy individuals (H1 to H7), their PBMCs were collected and used for the following analyses. The overall study design, schematic illustration of sample collection and processing are shown in Fig. 1A. Refer to the Material and Methods section for further details on the procedure. The personal information of the individuals is summarized in inset table (Fig. 1A, bottom, inset table).

**Figure 1.**
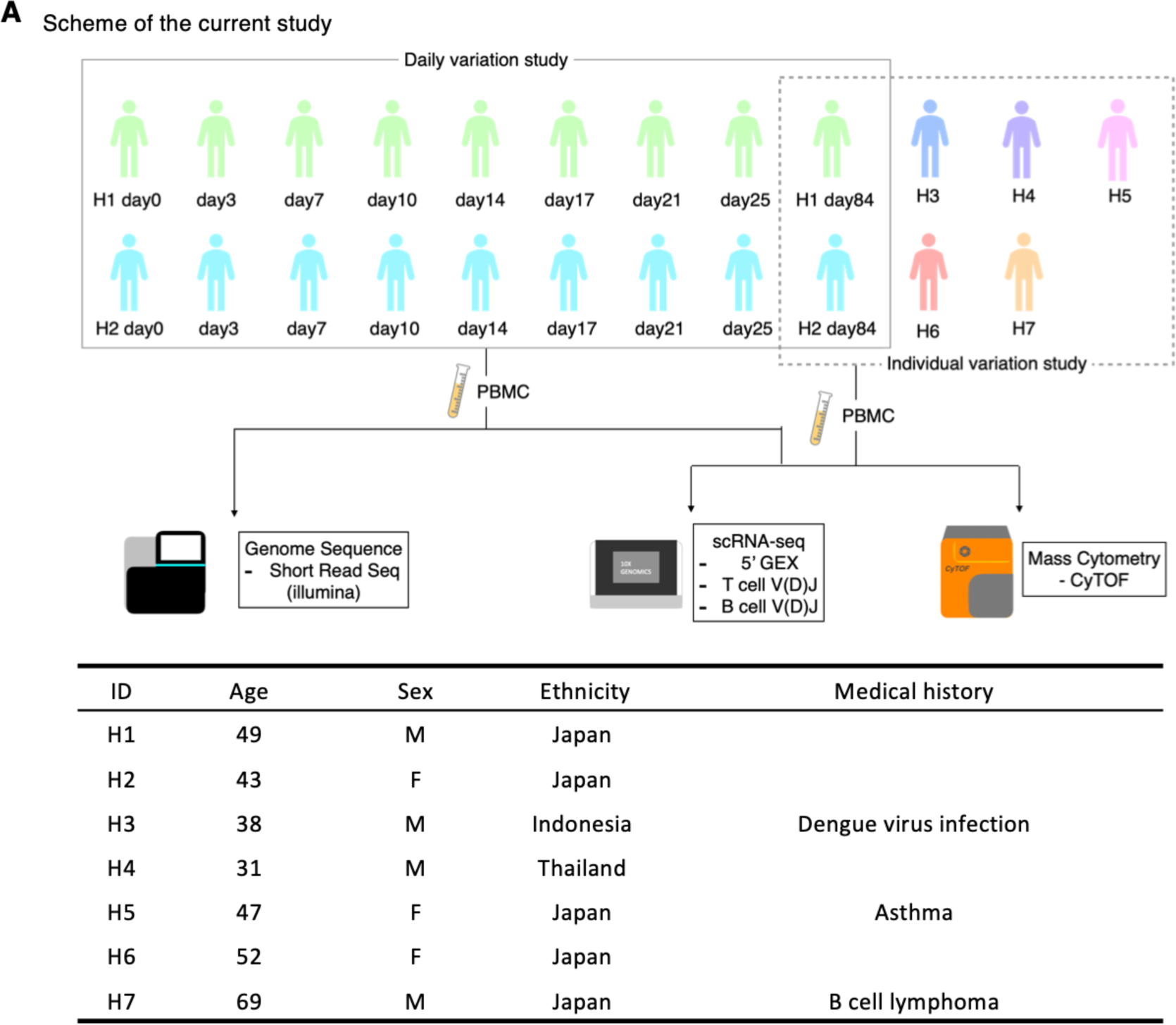
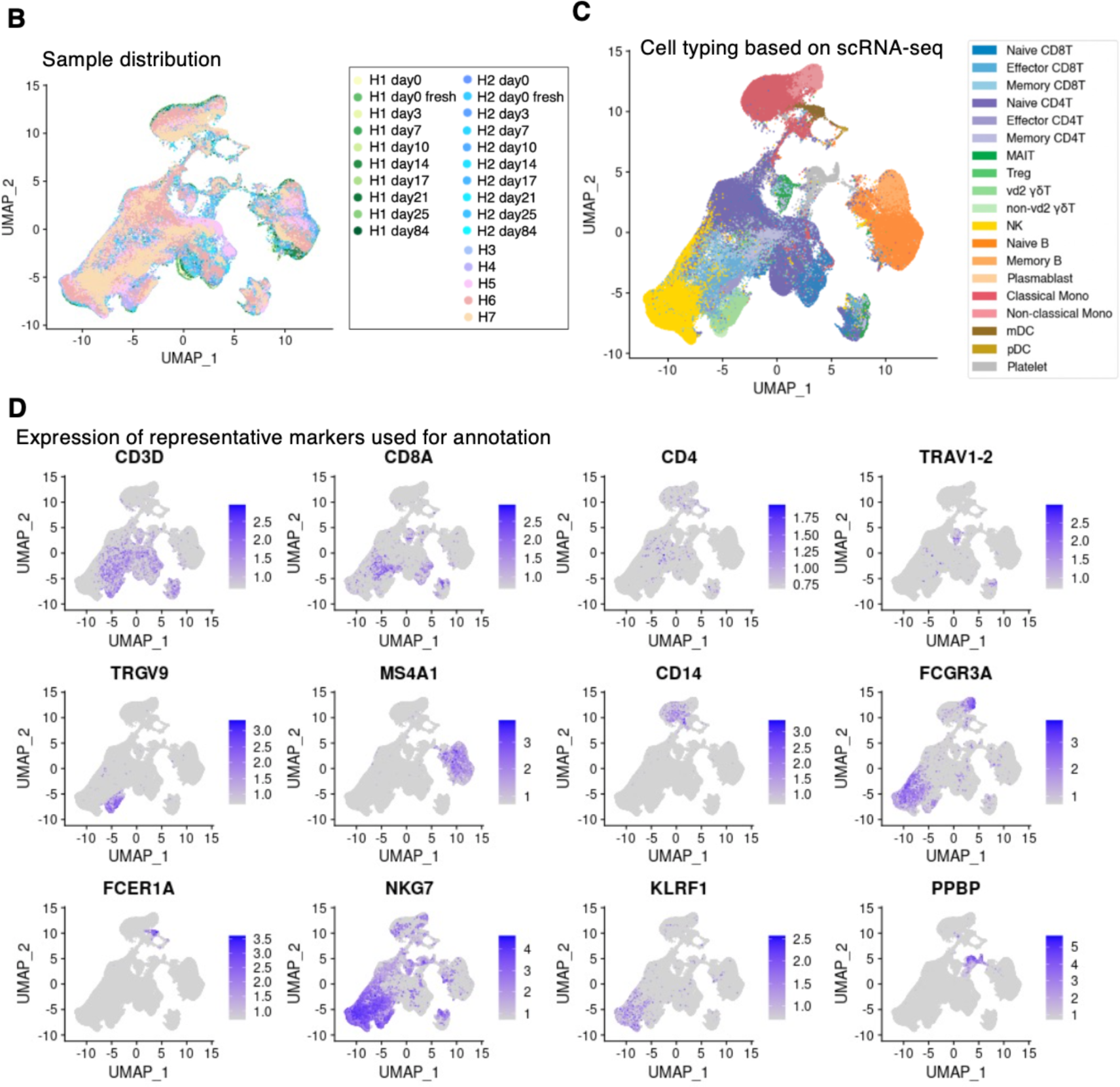
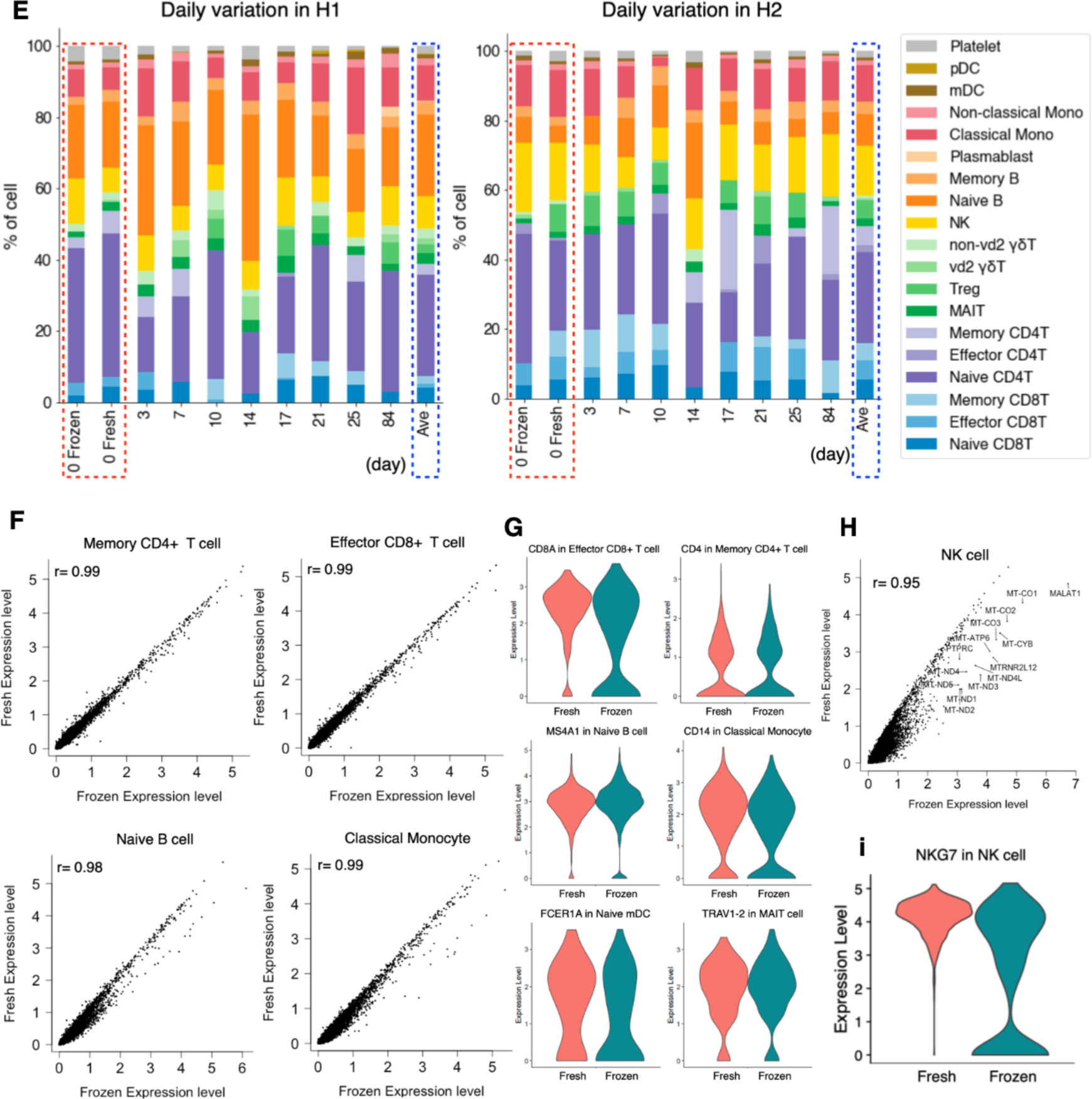
Characterization and evaluation of the scRNA-seq datasets. **(A)** Scheme and sample list of the present study. For the daily variation study, the multiple time-point samples were collected from H1 and H2 at the shown time-points. For the intra-individual variation study, PBMC samples were collected from seven participants, H1-H7. As illustrated the collected samples were subjected to the transcriptome, genomes, and proteome analyses (top). The medical history information about donors are shown in the table (bottom). **(B)** Evaluation of the sample distribution. We used the UMAP plot to confirm the existence of the batch effect. Each point shows a cell and is colored with 25 cases shown in the margin. **(C)** Cell type annotation. UMAP plot showing clusters colored by cell types. **(D)** Expression of representative markers for cell annotation. We used the following markers: CD3D, CD8A, and CD4 (T cell), TRAV1-2 (MAIT cell), TRGV9 (*γδ* T cell), MS4A1 (B cell), CD14 and FCGR3A (monocyte), NKG7, and KLRF1 (NK cell), PPBP (platelet). **(E)** Structure of PBMC at each time point of H1 (left) and H2 (right). The *x-axis* shows the day after first sampling, and the *y-axis* shows the percentage of each cell component. Bars with red dotted line show the data comparison of a fresh and frozen sample, and blue dotted line shows the average of each person. **(F)** Evaluation of the correlation between the fresh and frozen samples. We used H1 Day 0 fresh and H1 Day 0 frozen. The *x-axis* shows the expression level of the frozen sample, and the *y-axis* shows the expression level of the fresh sample. Correlation is shown in each plot. **(G)** Expression of indicated genes in each cell type of fresh and frozen samples. **(H)** Correlation of the gene expressions between the fresh and frozen NK cells. Mitochondrial genes are highlighted in the panel. Outlier genes are also shown in the plot. **i,** Expression levels of the NKG7 gene in fresh and frozen NK cells.

A droplet-based scRNA-seq (Chromium of 10X Genomics) was performed for all samples. Particularly for H1 and H2, PBMCs were sampled at nine-time points over months (Fig. 1A; Table S1). On average, 240,207,958 reads were obtained for a single sample. An average of 28,826 reads were assigned for a single cell as the 5’-end mRNA gene expression information (Table S2). Each sample was individually clustered and visualized by UMAP (Figs. 1B and 1C). Even without employing a batch-effect removing procedure, the images were mostly overlapped between individual experiments (Fig. 1B; for more details, see Table S2). Each cluster was annotated for a cell type using canonical cell markers (Fig. 1C and 1D). The cells belonging to each cell type were counted as the corresponding cell populations. The relative percentage of major cell types constituting the PBMCs was calculated for all the datasets (for statistics, see Tables S3 and S4).

First, we evaluated the reproducibility and reliability of the data obtained. At the same time, the effect of sample freezing was also evaluated. It is convenient to freeze samples after collection and keep them in a freezer until an appropriate time for library preparation. However, the exact effect sample freezing has on the PBMC transcriptome has not been fully evaluated. For this purpose, we prepared libraries from the same material to subject to two conditions: fresh (H1 Day 0 Fresh and H2 Day 0 Fresh) and frozen (H1 Day 0 frozen and H2 Day 0 frozen) (Fig. 1E, and red dotted line). Except for several specific particular cell types or a small group of genes in particular cell types that were excluded from the following analysis, there was no noticeable difference between the fresh and frozen sample in the total cell populations and gene expressions (Fig. 1F-1I; also note that some NK cells seemed damaged by the sample freezing, which were detected as the increased representation of mitochondoria genes; other low correlated genes are shown in Table S5). Therefore, we used frozen samples for further analyses. In the following analyses, we will describe some characteristic features between different individuals (see below). However, these distinct features were within the range of daily changes; therefore, each data was represented by nine independent experimental replicates. Collectively, we concluded that the collected data should be highly reproducible and reliable for the following analysis.

### Diversity of the scRNA-seq profiles between different individuals and different time points

When we examined the resulting scRNA-seq profiles (Fig. 1E, and Table S4), the annotated cell type composition roughly agreed with those previously estimated(Verhoeckx et al. 2015), that is, typically, lymphocytes (T cells, B cells, and NK cells) ranging 70-90 %, monocytes accounting for 10-20%, and DCs and other populations being rare. Within the lymphocyte population, cell types include CD3^+^ T cells (70-85 %), B cells (5-10 %), and NK cells (5-20 %). The CD3^+^ T cells consist of CD4^+^ T cells and CD8^+^ T cells in approximately 2:1 ratio.

Despite the overall concordance with previous estimates, cell compositions differed across individuals and sampling time points even within the same individual. At a glance, higher proportions of B cells were detected in H1 (Fig. 2A). On the other hand, CD8^+^ T cells and NK cells proportions were higher in H2 (Fig. 2A, Table S3). More specifically, naïve B cells and non-vd2 γδ T cells were highly represented in H1 (Fig.2A). Unlike usual T cells expressing α and β TCR chains, these non-vd2 γδ T cells do not necessarily require antigen representation via the MHC class I molecule for their activation(Weese et al. 2012), although their antigen recognition mechanism has not been fully characterized. These results suggested the possibility that the H1 immune landscape might be inclined to the humoral immune mechanism. On the other hand, the immune system in H2 can put a greater strain on the TCR-dependent response. Although some variations depended on the time points, these differences were characteristic to the individuals with statistical significance (p-values are shown in panels and legend; Fig. 2A).

**Figure 2.**
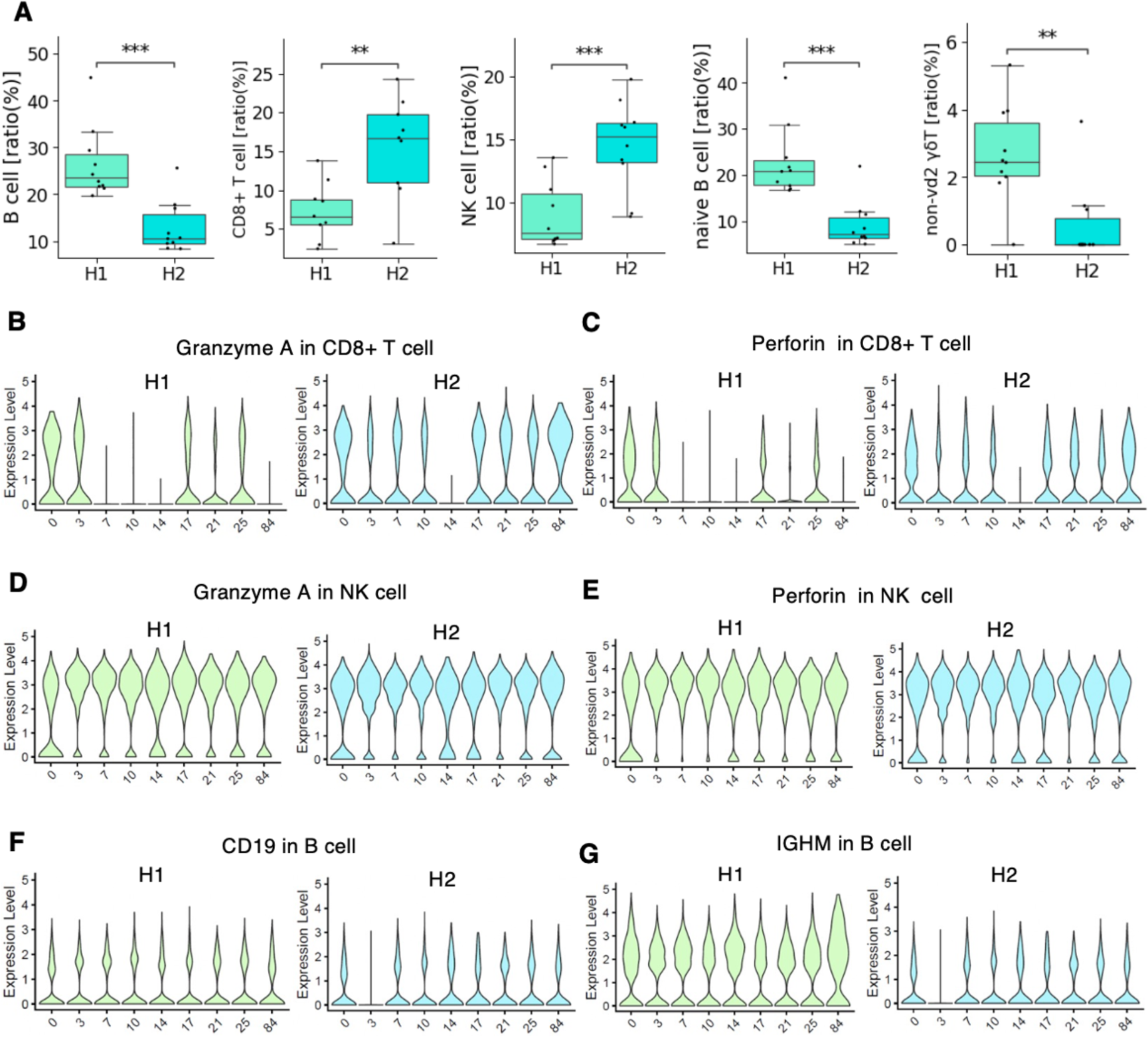
Daily diversity of PBMC profiling in H1 and H2. **(A)** Ratio of each cell type in H1 (pale green) and H2 (light blue). Boxplots include eight timepoints of each person of B cell, CD8^+^ T cell, NK cell, naïve B cell, non-vd2 gd T cell (left to right). p-value was calculated by t-test and shown as **: 1.00e-03 < p <= 1.00e-02, ***: 1.00e-04 < p <= 1.00e-03. **(B-G)** Expression level of representative genes in each cell type of H1 and H2; granzyme A expression of CD8^+^ T cell (**B**), perforin expression of CD8^+^ T cell (**C**), granzyme A expression of NK cell (**D**), perforin expression of NK cell (**E**), CD19 expression of B cell (**F**), and IGMH expression of B cell (**G**) in H1 and H2.

We also characterized the activation states by measuring gene expressions across different time points using the representative cell types in H1 and H2 in order to understand how active the immune cells in healthy subjects. For representative active markers of CD8^+^ T cells and NK cells, we found that their expression levels were almost similar between H1 and H2, in spite that some daily changes were observed. Figure 2B-2E exemplifies the case of Perforin (Osińska et al. 2014) and Granzyme A (Shi et al. 1992; Hayes et al. 1989). A similar observation was obtained for the activation state of B cells (Figs. 2F and 2G). These results suggested that the difference between H1 and H2 under healthy conditions was rather represented by the number of the corresponding cells, but not always the activation states of the individual cells.

### Immune cells diversity in seven individuals of varying backgrounds

To further assess the diversity of cell compositions across individuals, we compared the cell type proportions of the other seven individuals (Figs. 3A and 3B). The ninth (final) sample was taken as a representative for H1 and H2. As described above, H1 had a higher proportion of B cells than H2, and this trend even remained the most relevant among all samples (Figs. 3A and 3B). Particularly, the plasmablast population was far higher than the average of the other samples (2.8% and 0.15% for H1 and H2, respectively). On the other hand, H3 and H4 showed even higher frequency of non-vd2 γδ T cells than H1, suggesting that this feature is not totally unique to H1. Furthermore, H2 had high representations of memory CD4^+^ T cells, effector CD8^+^ T cells and MAIT cells in H3, and vd2 γδ T cells in H4 (Fig. 3B, Table S3). All individuals showed, in part similar, but a wide variety of unique features.

**Figure 3.**
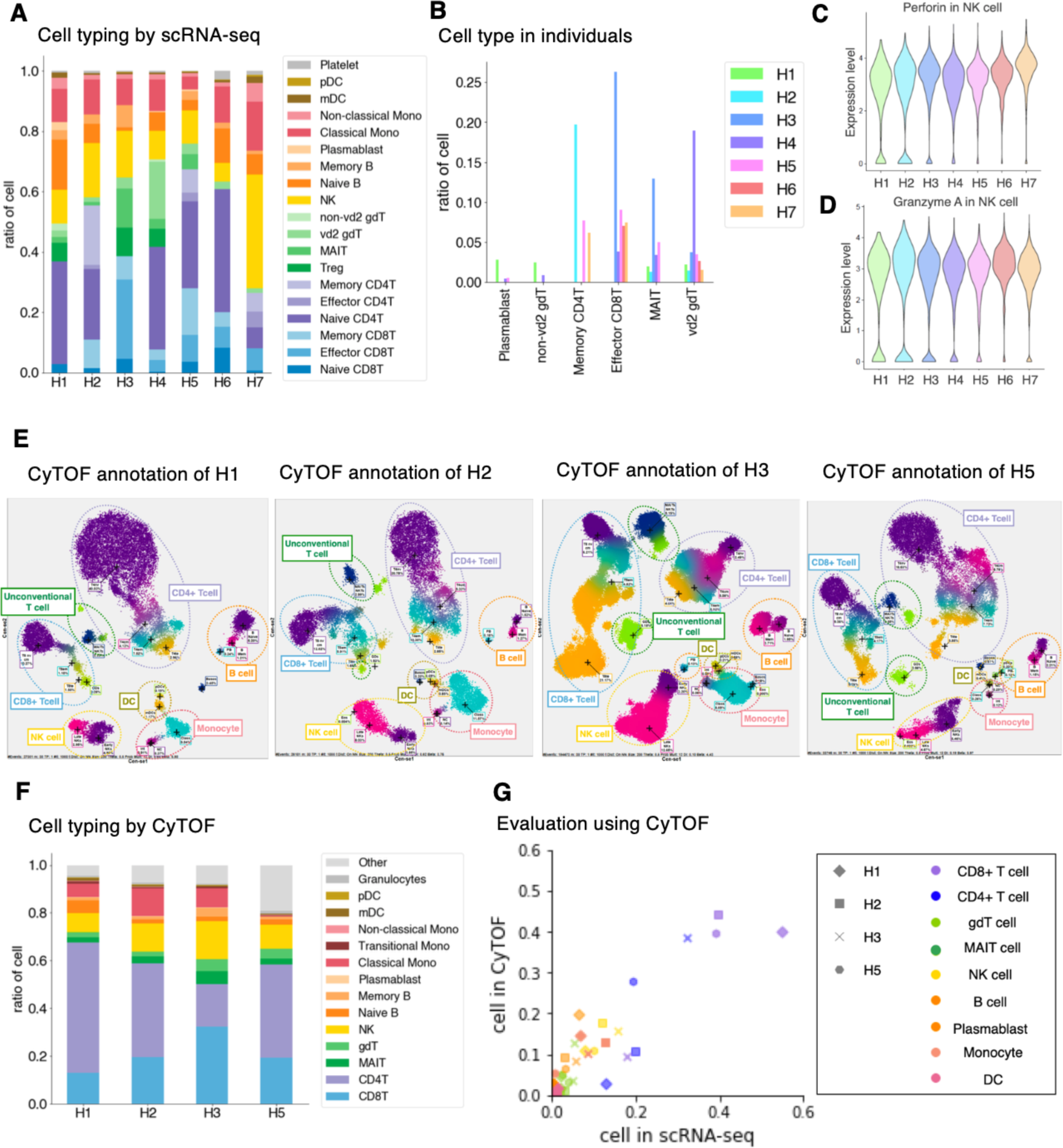
Diversity of PBMC profiling in seven individuals. **(A)** Structure of PBMC in seven individuals. Cells are annotated based on the gene expressions in the dataset analyzed by scRNA-seq. The *x-axis* shows the individuals, and the y-axis shows the ratio of each cell component. Color legends are shown in the margin. **(B)** Individual variance of cell types. The *x-axis* shows the focusing cell type, and the *y-axis* shows the ratio of cells in individuals. **(C and D)** Gene expression level in seven individuals. Gene and cell type of interest are shown at the top of the graph. **(E)** Cell typing using CyTOF of H1, H2, H3, and H5 (from left to right). **(F)** Structure of PBMC based on CyTOF. The *x-axis* shows the individuals, and the y-axis shows the ratio of each cell type. Color annotations are shown in the margin. **(G)** Evaluation analysis using CyTOF. Scatterplot showing the correlation of the ratio of cells annotated by scRNA-seq (*x-axis*) and by CyTOF (*y-axis*). Markers are shaped depending on each individual and colored by cell types shown in the margin. For H1 and H2 used in Figure 3, we selected final time-point as representative samples.

Among them, H7 showed a unique profile (Fig. 3A). The cellular population and the gene expression profiles of individual cells suggested that NK cells are in the active state in this individual (Figs. 3C and 3D). This individual is an older adult and has experienced malignant B cell lymphoma (Fig. 1A, bottom, inset table). The cytotoxicity of NK cells has a high anti-tumor potential(Vivier et al. 2008). NK cells are often suppressed in blood cancer patients when the disease is in a malignant stage. However, as patients recover, the reactive population of NK cells increases, resulting in disease remission(De Kouchkovsky and Abdul-Hay 2016). Although more than five years have passed since the complete elimination of malignant B cells by successful R-CHOP chemotherapy, the remaining large proportion of NK cells in H7 may have expanded during therapy. A recent study reported that prolonged expansion of clonal NK cells occasionally occurs after recovery. In fact, a sustained expansion of NK cells may suggest clonal expansion beacause of response to any chronic stimutlation(Adams et al. 2020) or or acqusition of somatic mutations(Olson et al. 2021). Collectively, the results suggest that healthy individuals hold a prominent baseline immunological diversity.

Before further exploring the observed difference, we considered the validation using other methods to validate whether the observed diversity should correctly represent the diversity between individuals. For this purpose, Cytometry of Time-Of-Flight (CyTOF), analysis was employed. This method utilizes the mass cytometer HeliosTM, using heavy metal isotope-tagged antibodies to detect PBMCs proteins at the single-cell resolution(Spitzer and Nolan 2016). Four samples (H1, H2, H3, and H5) were subjected to the analysis (Figs. 3E and 3F). Comparing the transcriptome and proteome datasets showed that the detected cellular compositions were roughly equivalent regardless of the analytical methods (Fig. 3G). It is true that, the transcriptome data gave a larger inclination than the proteome data for some cell types, while the opposite was observed for other cell types (see Table S6 for details). Probably these observations were because mRNA and protein levels are not strictly equal. Nevertheless, we found a high correlation between transcriptome and proteome data in almost all cases. Thus, the observed diversities of immune cell profiles are validated from this viewpoint.

### Time-lapse changes of the immune landscapes in T cell populations

We attempted to further characterize the diversity of immune cell responses by considering the VDJ regions of TCR or their “clonotypes”. Using the Chromium platform, the VDJ-seq libraries were constructed from the intermediate products of the library construction for the scRNA-seq (see Table S7 for the sequencing statistics). Since the cell barcodes were shared between the VDJ-seq and scRNA-seq libraries from the same sample, we could associate the observed VDJ information with the transcriptome information of its expressing T cell for each cell. Similarly, in the case of scRNA-seq, for H1 and H2, data was collected from nine points for H1 and H2 over a month (H1 Day 0 to H1 Day 84, and H2 Day 0 to H2 Day 84, Table S1).

For H1 and H2, even the ten most frequent clonotypes claimed a small proportion of the overall annotated cell population. Furthermore, the ten most frequent clonotypes were unique to each individual, and no explicit overlap was observed (Fig. 4A). Nevertheless, some features in the pattern of the compositions and their changes were commonly observed between H1 and H2 (Fig. 4B; see Table S8 for more details). We examined and found that the clonotypes that were unique within the same individual over different time points (“sporadic” clonotypes) were mostly from naive T cells, probably representing a unique repertoire of unstimulated T cells in the individual (Figs. 4D-4F). On the other hand, as for the clonotypes detected from several time points, effector and memory CD8^+^ T cells were dominant (Figs. 4D-4F). Of note, from those “sustained” T populations, MAIT cells accounted for a significant population (Fig. 4F).

**Figure 4.**
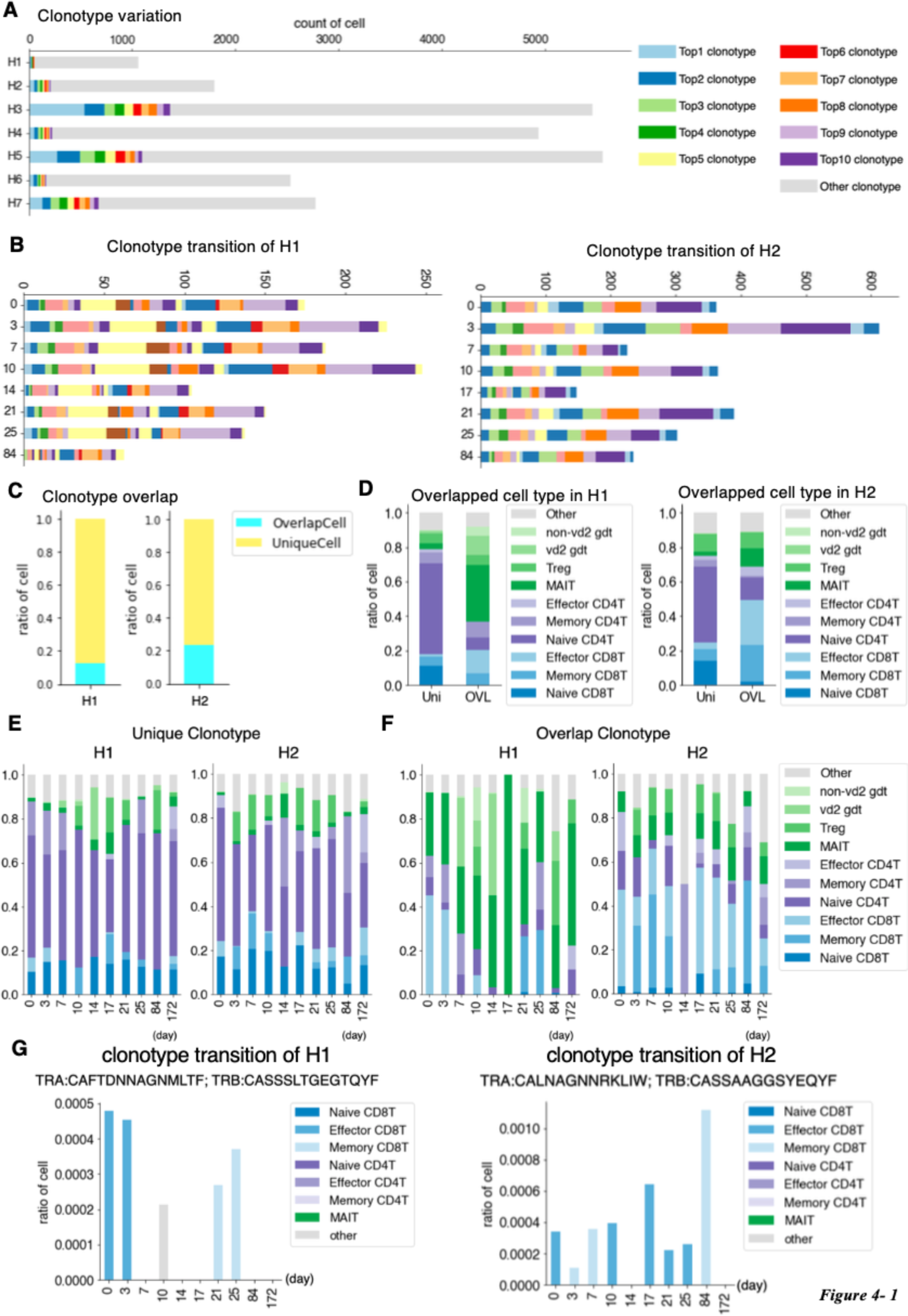
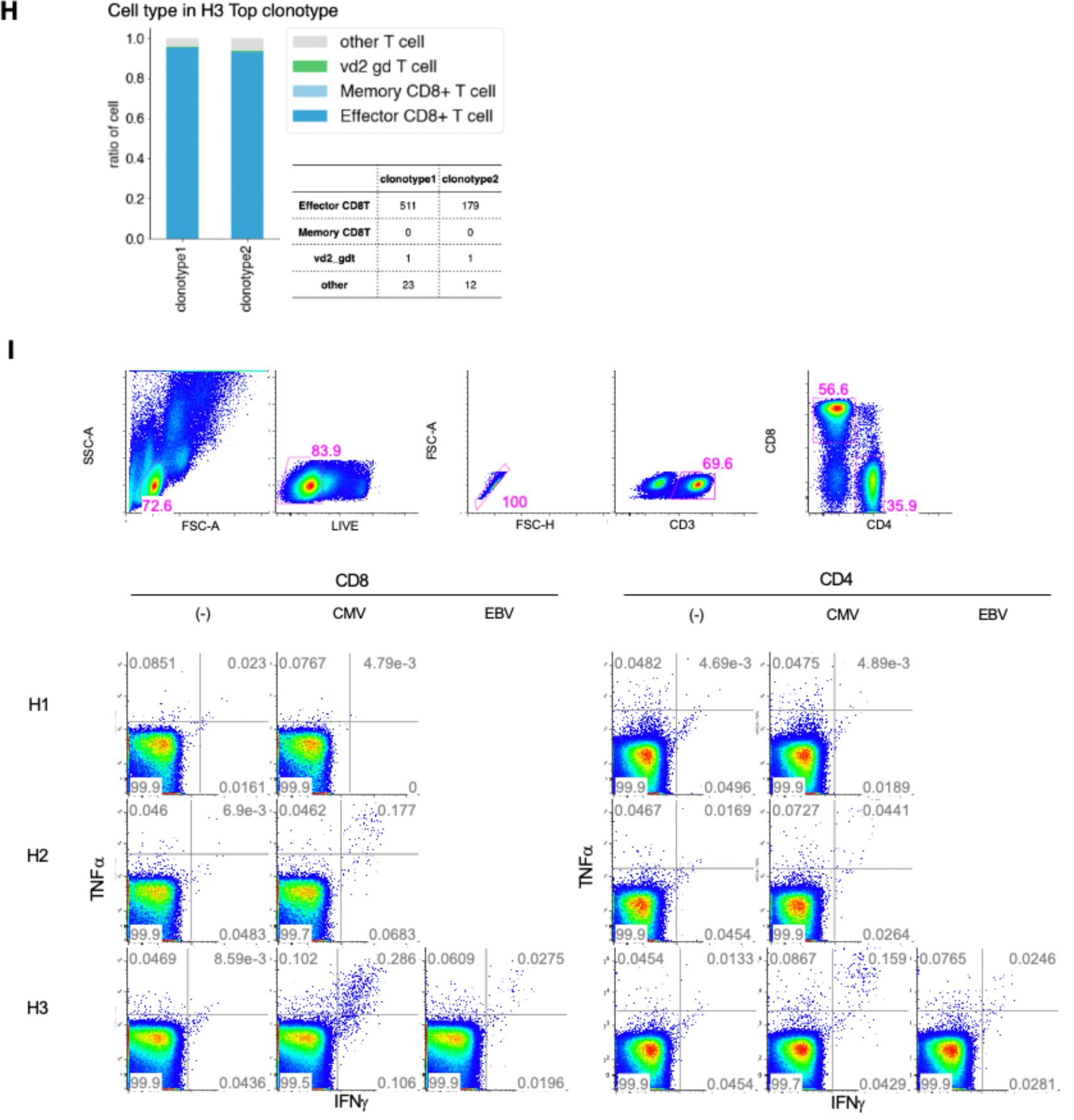
TCR individual and daily variation. **(A**) Clonotype divergence between seven individuals. The *x-axis* shows the number of T cells and the *y-axis* shows the individual. Cell with top 1-top 10 clonotypes are plotted as specific colors shown in margins. Top clonotypes shown are not common between individuals. **(B)** Clonotype divergence of nine timepoints of H1 (left) and H2 (right). Barplot shows clonotypes ranked as top 1-top 10 at any point in time. Note that color in H1 and H2 is not common. **c,** Barplot showing the ratio of T cells with unique TCR (light blue) and overlapped TCR (yellow) in H1 (left) and H2 (right). **(D)** Barplot showing the ratio of the detailed T cell type in unique (Uni) and overlapped (OVL) of H1 average (left) and H2 average (right). **(E and F)**, Barplot showing ratio of detailed T cell type in unique (**E**) and overlapped (**F**) of H1 (left) and H2 nine time points (right). **(G)** Barplot showing the ratio of T cells at each time point with specific clonotype TCR of H1 (left) and H2 (right). Information about the TCR is shown at the top of each graph. **(H)** Cell lineage of H3 top1 and top2 clonotype. **(I)** ICS analysis. top panel: cell type separation is shown for H3 as a representative (top); ICS analysis using H1 (top), H2 (middle) and H3 (bottom) under no stimulation as control (left) and peptide stimulation of CMV (middle), EBV (right) in CD8^+^ T cell (right) and CD4^+^ T cell (left).

Interestingly, we could trace the time-lapse transition of their expressing T cell for some of those sustained clonotypes (Fig. 4F). For example, for a particular clonotype, as shown in Fig. 4G, its expressing T cells were effector CD8^+^ T cells. This proportion decreased within a week and memory CD8^+^ T cell appered on Day 21. This individual could have been infected with a pathogen in his/her self-presumed healthy state (Fig. 4G, left). Similar situation was also confirmed in H2 (Fig. 4G, right). Accordingly, the activation of T cell populations may constantly occur even in the “healthy” condition. We further searched from H1 and H2 and identified a total of 85 and 209 clonotypes increased and decreased during this time-frame. Immune cells may undergo constant changes, responding to environmental epitopes, and such responses may have shaped the unique immune landscape of the individual over long years.

### Diversity of TCRs and searching for their possible epitopes

We conducted a similar TCR analysis for the other individuals. Despite the reduced data points for other samples, similar trends were also observed for H4 and H6, although their exact VDJ sequences were, again, unique to the individuals (Fig. 4A). Of note, in H3, the most and the second-most frequent clonotype claimed were clonotype 1 (13.4%) and clonotype 2 (4.8%), respectively (Fig. 4A). We carefully ruled out the possibility that these were derived from PCR and other artifacts by manually inspecting the correct assignment of cell barcode and unique molecular index. Particularly for this clonotype, we dissolved their TCR states by utilizing the scRNA-seq information. We found that clonotype 1 and 2 were mostly for effector CD8^+^ T cells, suggesting that some asymptomatic infection events are on-going (Fig. 4H).

As for H3, this individual is originally from a suburb in Indonesia. Considering the country of his/her upbringing, we postulated that this individual may have frequently experienced the infections of CMV, EBV and other common pathogens. We conducted the intracellular cytokine staining assay (ICS assay) for H1, H2 and H3 samples (Fig. 4I). We foumd that H3 showed the highest response for the CMV stimulation. Similar responses were found for EBV (Fig. 4I). In H3, the immune system generally may remain alerted, which could be a common trait for individuals originally from developing countries. Supportingly, when we attempted to infer potential epitopes for the clonotypes detected in H1-H3, using the deduced amino acid sequences of the CDR3 region, which is the docking platform for the epitopes, for the bioinformatics prediction pipeline and epitope databases TCRex(Gielis et al. 2019) and VDJdb(Shugay et al. 2018), we found that cytomegalovirus (CMV) amd Epstain-Barr virus (EBV) appeared to be potential candidates for the TCRs more frequently in H3 (Tables S9).

### Diversity of BCRs

We conducted a similar analysis for BCRs (sequence statistics are shown in Table S7). Even to a lesser extent than the TCRs, the BCR clonotypes did not overlap within the same individual over time, particularly for the usage of the immunoglobulin heavy chain (99% were uniquely observed; Fig. 5A). The major unique clonotypes were mostly for IgH-M in H1, suggesting that there is constant activation of B cells for possible novel antigens in this individual (Fig. 5B). On the other hand, IgH-G was more relevant within a minor population of overlapping clonotypes at different time points, and possibly represented the sustained activation of the corresponding clonotypes. A similar trend was observed for H2 (Fig. 5B), but to a lesser extent than H1. We further examined the overall entropy of the BCRs. The distribution of the Shannon index showed that H1 had the higher entropy than H2 and the other individuals (Fig. 5C), although daily changes were observed in this aspect (Fig. 5D). As well as TCRs, diversity of BCR in naïve B cells are considered to be higher than that of memory B cells, which have already experienced clonal expansion by specific antigen stimulation. The higher entropy of BCRs in H1 may reflect higher frequency of naïve B cells. When we analyzed the frequency of the variant regions, we found that the H1 showed a focused use of particular variant types, although their precise clonotypes were diverse (Fig. 5E). These results showed that BCR profiles also vary between individuals. They also collectively indicated, again, that B cell-mediated immune responses are prominent in H1.

**Figure 5.**
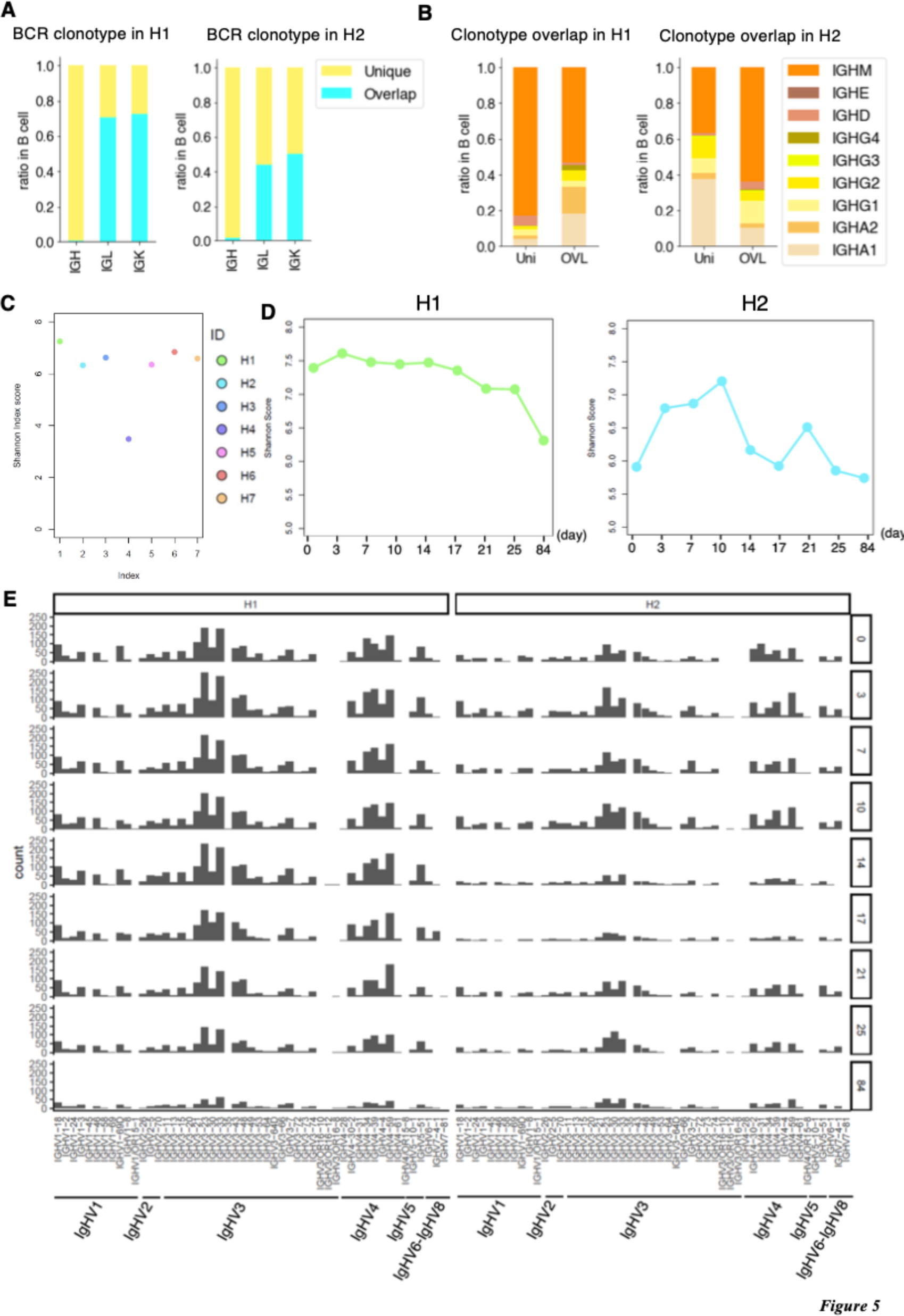
Individual variation and Daily variation of BCR. **(A)** BCR clonotype divergence in H1 (left) and H2 (right). Barplot showing the ratio of overlapped (paleblue) and unique (light yellow) clonotype of IgH, IgL, and IgK. **(B)** Ratio of each clonotype of unique(left) and overlap(right) in H1 and H2. **(C)** Shannon index score variation of BCR in H1-H7. Color legend is shown in the margin. For H1 and H2, we used the average score of Day 0 to Day 84. **(D)** Daily variation of Shannon index score of BCR daily variation in H1 (left) and H2 (right). The *x-axis* shows the time point, and the *y-axis* shows Shannon’s score. **(E)** Variation of V gene in BCR from Day 0 to Day 84 (top to bottom) in H1 (left) and H2 (right).

### Influence of the vaccination on the immune cell profiles

We considered vaccination an ideal usual life event to further characterize the personal immune landscapes and their changes. During vaccination, the exact antigen is defined, and the exposure time is known. We first collected PBMC samples from H1 and H2 before and after influenza vaccination (antigen was for the 2020, see Methods). Relevant antibodies titers were confirmed by the antibody quantification method (Table S10). Samples were collected on Day - 1, 1, 3, 7, and 28 of vaccination and subjected to similar scRNA-seq and VDJ seq analyses for TCRs and BCRs (Fig. 6A and see Tables S2 and S7 for the sequencing statistics).

**Figure 6.**
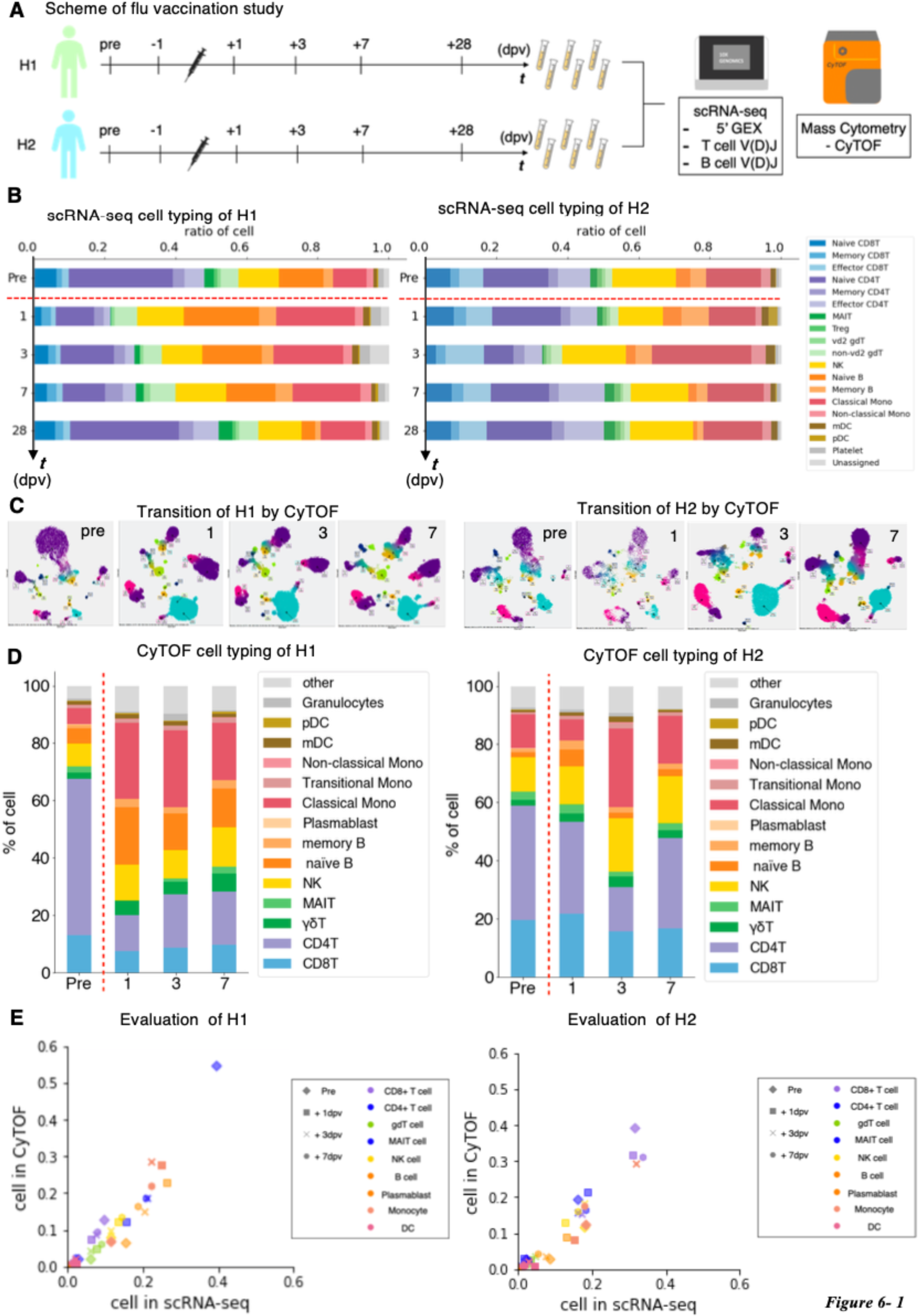
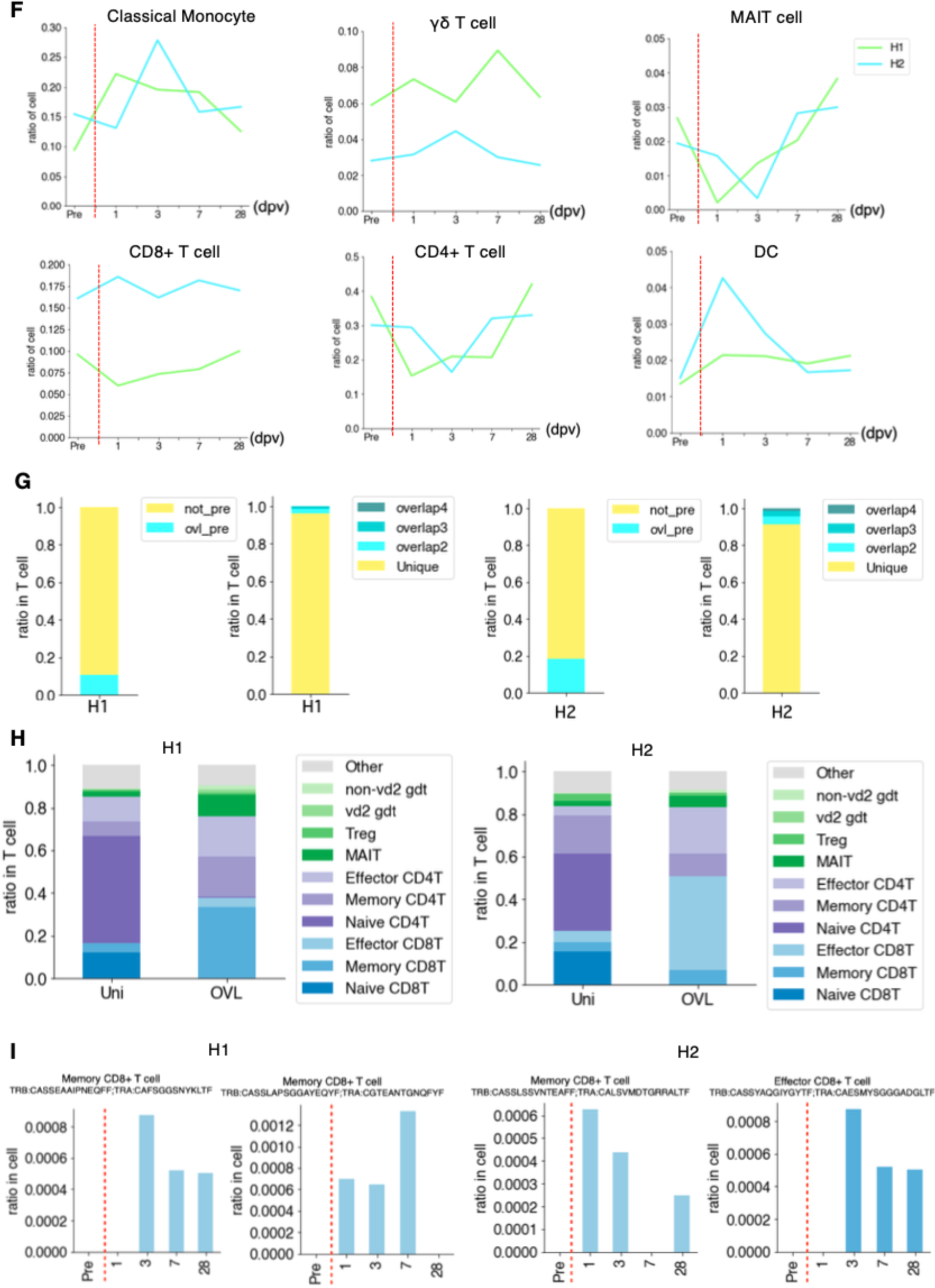
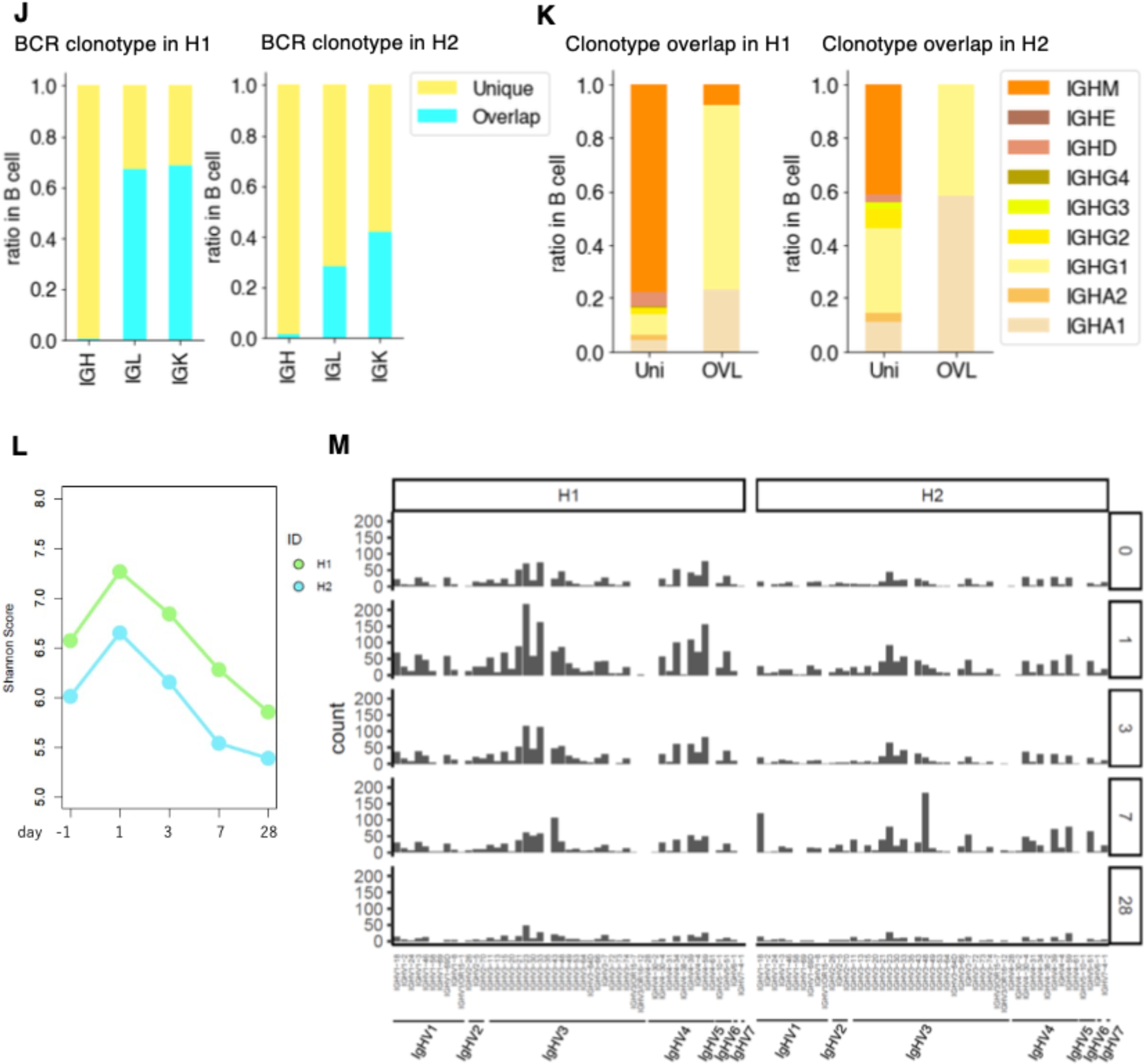
Perturbation of the immune cell gene expression profiles depending on influenza vaccination. **(A)** Influenza vaccination study scheme. We collected blood samples from H1 and H2 one day before vaccination (-1 dpv, day post vaccination), 1, 3, 7, and 28 dpv. We analyzed PBMCs transcriptome, V(D)J of BCR and TCR in single-cell level using 10x Genomics, and mass cytometry in single-cell level using Fluidigm CyTOF. **(B)** Barplot showing the transition of cell components before and after influenza vaccination of H1 (left) and H2 (right) analyzed by scRNA-seq. **c**, Cen’s plot showing the transition of cell components analyzed by CyTOF of H1 (left) and H2 (right) in pre, 1, 3, 7 dpv. **(D)** Barplot showing the transition of cell components analyzed by CyTOF of H1 (left) and H2 (right) in pre, 1, 3, 7 dpv. Color annotations are shown in the margin. **(E)** Scatterplot showing the correlation of the ratio of cells annotated by scRNA-seq (*x-axis*) and by CyTOF (*y-axis*). Markers are shaped depending on timepoint and colored by the cell types shown in margin. (**F)** Transition of the ratio of cell type shown in lineplot. The *x-axis* shows day post vaccination and the *y-axis* shows ratio of cell in each person. (**G)** Barplot showing the ratio of T cells with unique TCR (light blue) and overlapped TCR (yellow) in H1 (left) and H2 (right). Clonotype exit in pre-vaccination is annotated as “ovl pre” and in only after vaccination annotated as “not pre”. Clonotype of “not pre” further grouped into Unique, only exit at one timepoint, and overlap (ovl). Number after ovl shows number of apprerance. (**H)** Barplot showing the ratio of the detailed T cell type in unique and overlapped of H1 average (left) and H2 average (right). (**I)** Barplot showing the ratio of T cells with specific clonotype TCR of H1(left) and H2 (right) in each time point. The *x-axis* shows day post vaccination, and the *y-axis* shows the ratio of specific T cells. Information about the TCR clonotype is shown at the top of each graph. **(J)** BCR Clonotype divergence in H1. Barplot showing the ratio of overlapped (pale blue) and unique (light yellow) clonotype of IgH, IgL, and IgK. **(K)** Ratio of each clonotype of unique(left) and overlap(right) in H1 and H2. **(L)** Shannon index of H1 and H2 BCR. **(M)** Diversity of the gene in BCR from -1 dpv to 28 dpv (top to bottom) in H1 (left) and H2 (right).

Again, diverse immune cell profiles between different individuals and time points were found for this time-lapse dataset. Significant differences were observed in response to the vaccination between H1 and H2 (Fig. 6B). The distinct responses appeared, representing their original immune landscapes (see below). Nevertheless, several common features were observed, generally consistent with previous knowledge describing general features of immune cell responses. For example, expansion of monocytes, primarily CD14^+^ classical monocytes, were detected as the primary responder of the stimulation immediately after vaccination. This induction was followed by a temporary reduction of naïve B cells in PBMC (Fig. 6B). CD4^+^ T cells and γδ T cells were also temporarily reduced for T cell populations, while CD8^+^ T cells, NK cells, and DCs cells retained their original population sizes. For these profiles, we also conducted the validation analysis using CyTOF and confirmed the robust representation of the observed results (Fig. 6C-6E). Of note, these initial responses were recovered to their original levels by Day 28 post-vaccination, when the immune responses are estimated to be complete and memory cells established (Fig. 6B).

Although the above responses applied to H1 and H2 in general, several unique features appread to depend on the individual. For example, in H1, the population of MAIT cells were particularly reduced from PBMC at the initial response from 2.7% to 0.2% in scRNA-seq (2.6% to 0.2% in CyTOF) (Fig. 6F, top right). In addition, the response of CD4^+^ T cells was more pronounced than that of H2 (Fig. 6F, bottom center), while the population of CD8^+^ T cells was larger at all times (Fig. 6F, bottom right), perhaps recapturing the dominant humoral responses in H1 (Fig. 6B).

We inspected the changes of TCR clonotypes in response to vaccination (Fig. 6G and 6H, Figure S1). The TCR repertoire and clonotypes detected before vaccination were removed to focus on the specific response to the vaccination. A total of 10 VDJ sequence datasets were used for the subtraction for each individual (see Tables S7 for the statistics).

For the remaining clonotypes obtained, we attempted to identify the clonotypes showing dynamic changes in response to the vaccination (Fig. 6I). Again, we observed that some clonotypes sporadically appeared in a particular time point; others were persistent. We compared the T cell populations between those sporadic and persistent populations. We found that the CD4^+^ and CD8^+^ naïve T cells were characteristic in the sporadic population, suggesting that these could be the clonotypes that were not removed by the subtraction. On the other hand, CD4^+^ and CD8^+^ memory T cells as well as MAIT cells were more relevant in the persistent population, suggesting that the clonotypes firstly induced by vaccination were enriched in this population. Some examples are shown for the clontypes which showed dynamic changes (Fig. 6I), as the candidate T cells first induced in response to vaccination. At least, a total of 16 and 53 such clonotypes were detected, in H1 and H2, respecstivery.

As for the BCR repertoire, we did not find any relevant overlapping clonotypes as shown in the above analysis (Figs. 6J and 6K). When we examined their complexity (Fig. 6L), we found that the entropy score increased on Day 3 as an immediate response of BCRs. This induction recovered to the original level gradually by Day 28. To different extent, this trend was commonly observed for both H1 and H2 (Fig. 6L). The variant analysis also showed that the particular variants were induced on day three (Fig. 6M). These results indicated that the B cell system was also responding to vaccination, again to varying degrees in different individuals.

### Responses of TCR clonotypes in response to the SARS-CoV-2 vaccinations

We conducted a similar analysis for vaccination against SARS-CoV-2 (see Tables S2 and S7 for the statistics). This time, the mRNA vaccine of BNT162b2 produced by Pfizer-BioNTech was considered. PBMC samples from H1, H6, and H7 individuals were used for analysis (Fig. 7A). Relevant increases in the antibody titers were measured by standard antibody quantification (Figs. 7B and 7C, Table S10).

**Figure 7.**
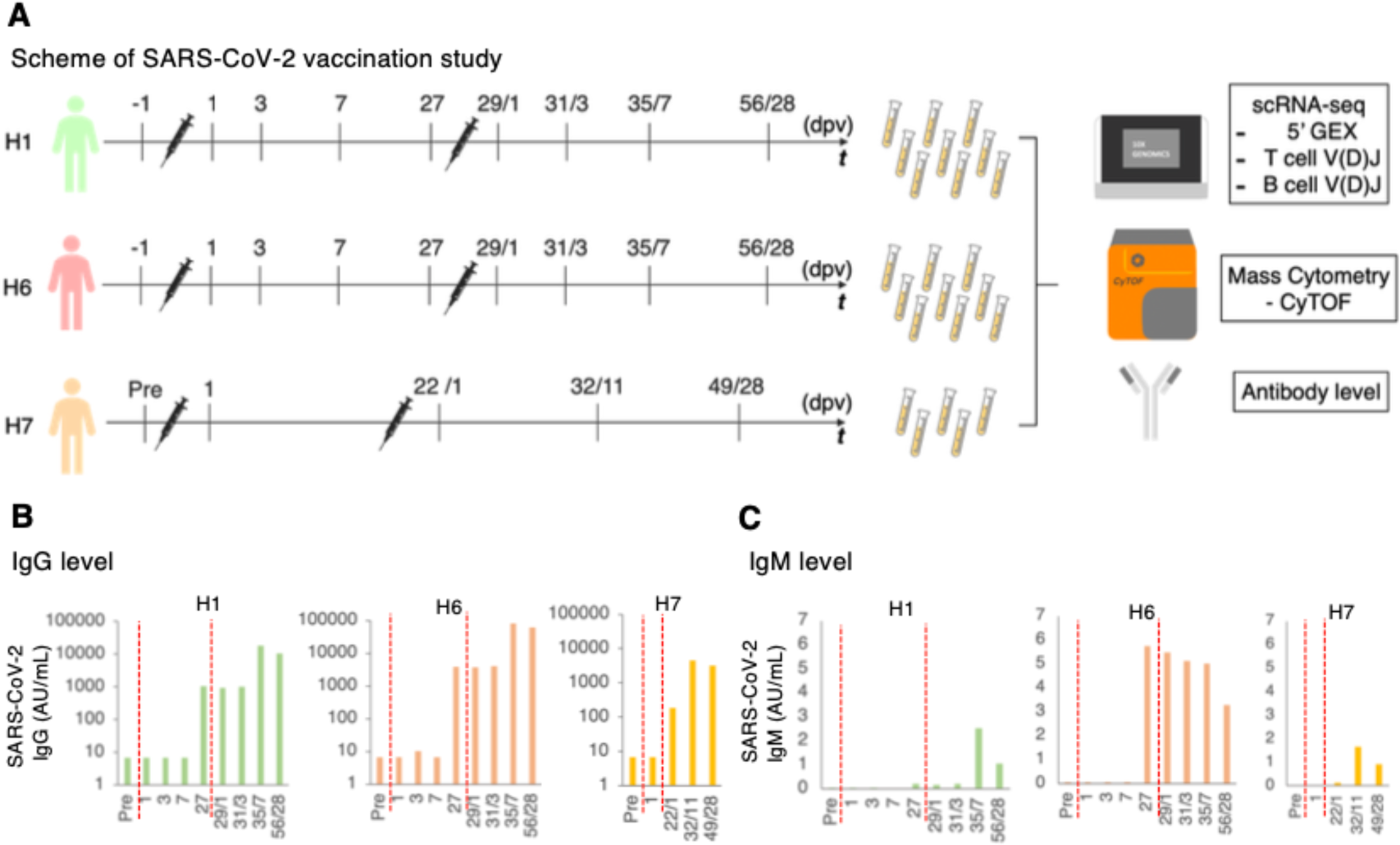
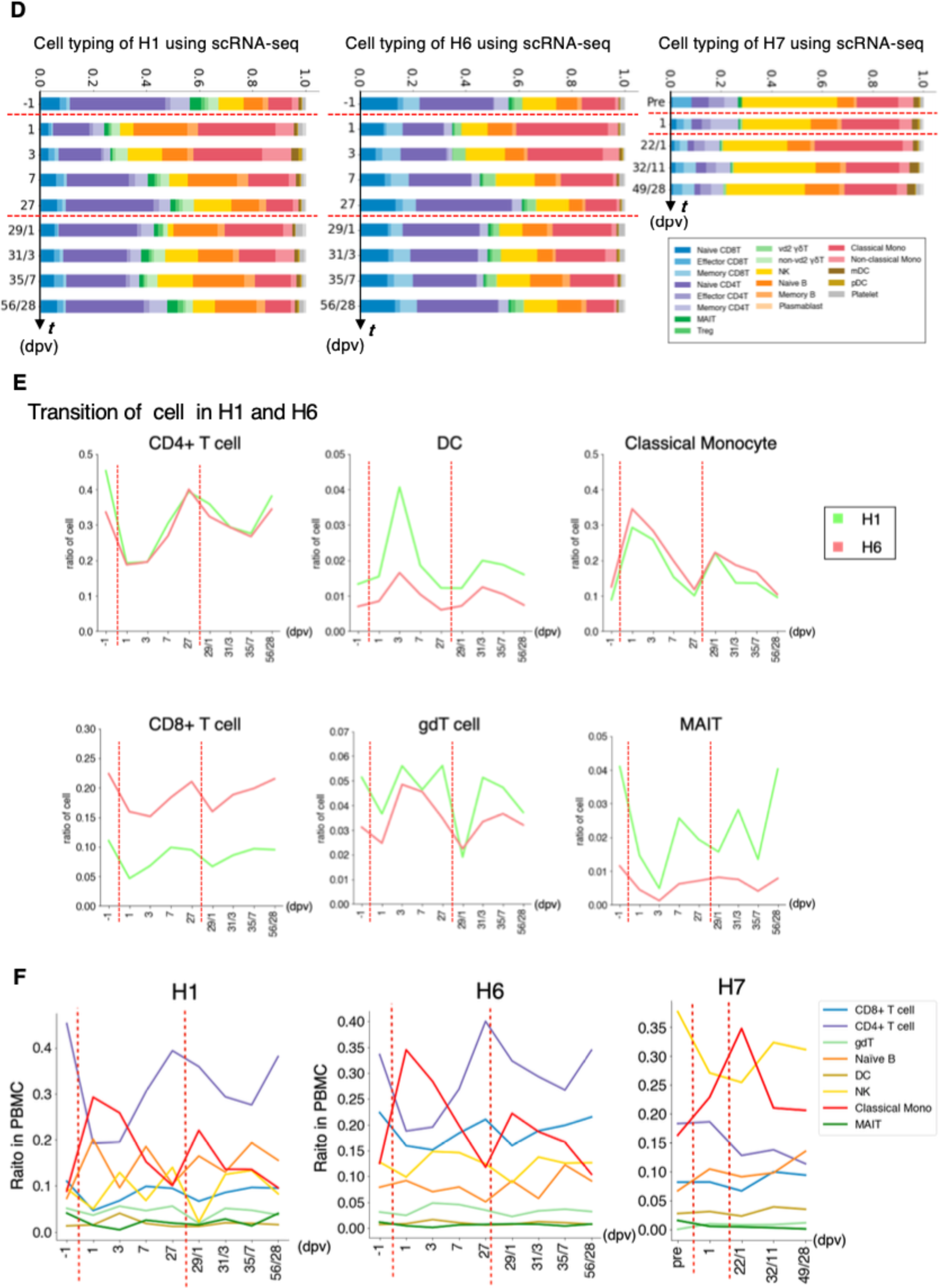
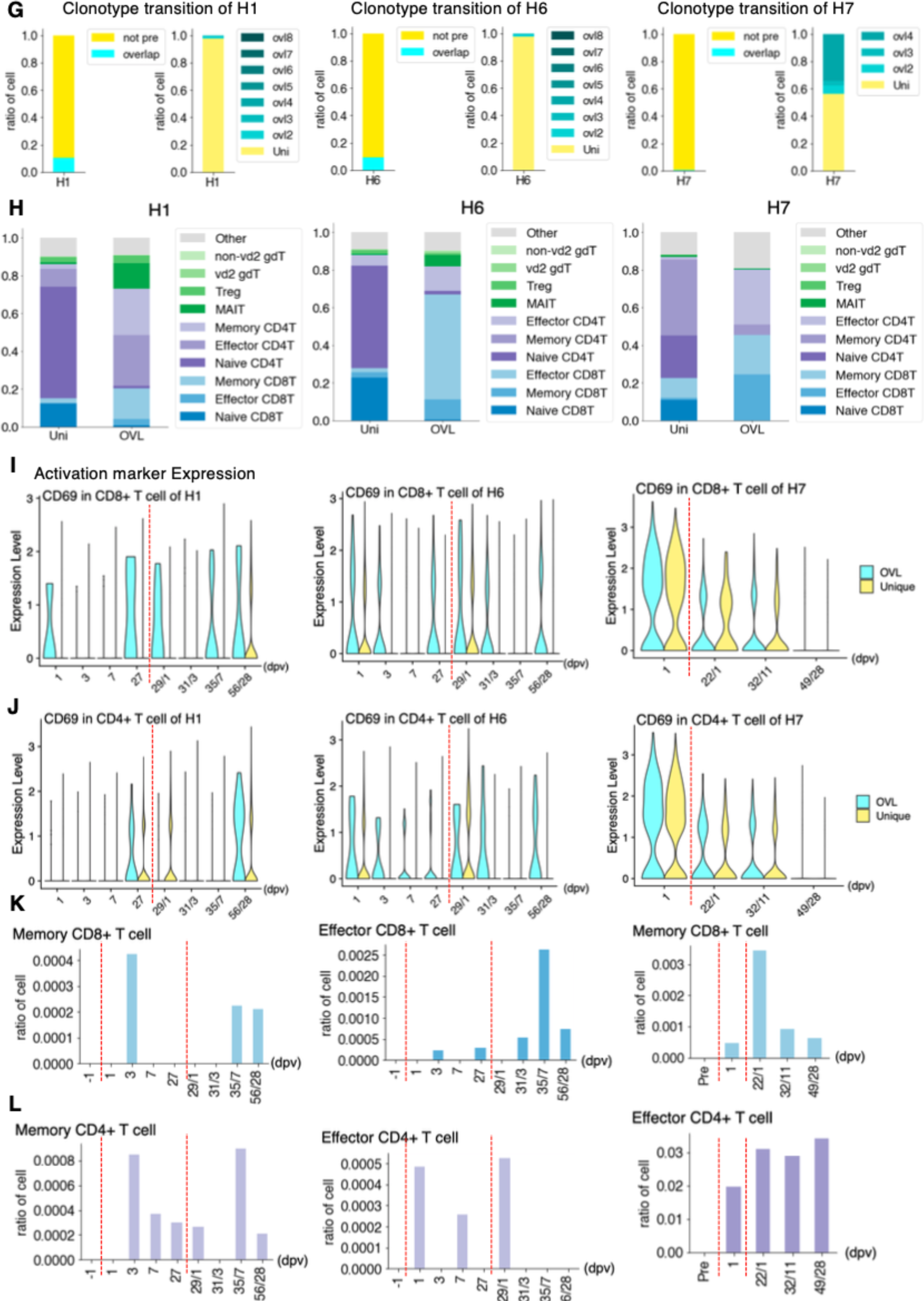
Perturbation of the immune cell gene expression profiles depending on SARS-CoV-2 vaccination. **(A)** SARS-CoV-2 vaccination study scheme. Blood samples of H1, H6, and H7 were collected before the vaccination (one day before vaccination, -1 dpv), post-first vaccination, and post-second vaccination. We analyzed PBMCs transcriptome, V(D)J of BCR and TCR at the single-cell level using 10x Genomics, proteomics in single-cell level using Fluidigm CyTOF and anti-SARS-CoV-2 virus antibody level. (**B and C)** Transition of anti-SARS-CoV-2 antibody levels. Barplot shows antibody level at each time point. The *x-axis* shows the day after the first vaccination, and the *y-axis* shows the level of IgG (AU/mL) **(B)** and IgM (AU/mL) **(C)** in H1 (green, left), H2 (coral, middle) and H7 (orange, right). **(D)**, Barplot shows the transition of cell components before and after SARS-CoV-2 vaccination based on scRNA-seq datasets of H1 (left), H6 (middle), and H7 (right). **(E and F)** Transition of cell type population before and after SARS-CoV-2 vaccination in H1 and H6 **(E)** and H1, H6 and H7 **(F)**. **(G)** Transition of T cell with unique clonotypes and overlapped clonotypes of H1 (left), H6 (middle) and H7 (right). Clonotypes exit in pre-vaccination is annotated as “overlap” and in only after vaccination annotated as “not pre”. Clonotypes of “not pre” were further grouped into Unique, as the ones which only exited at one timepoint, and the ones which overlapped (ovl). Number after “ovl” represnts the order of the appearance. **(H)**, Barplot shows T cell component transitions of H1 (left), H6 (middle) and H7 (right). **(I and J)** Expression level of the activation marker of T cell with a unique clonotype and overlapped clonotypes in CD8^+^ T cell **(I)** and CD4^+^ T cell **(J)**. The *x-axis* shows the day after the first vaccination, and the *y-axis* shows the CD69 gene expression. **(K and L)** Barplot showing the ratio of T cells with specific TCR in SARS-CoV-2 vaccination in H1 (left), H6 (middle), and H7 (right). The *x-axis* shows the day post-vaccination, and the *y-axis* shows the ratio of the specific T cells. Information about the TCR clonotype is shown at the top of each graph.

Similar features with influenza vaccination were observed here. These responses were also generally consistent with the results of a recently published paper(Ewer et al. 2021). These features include the immediate induction of monocytes and eventual restoration of the immune cell states (Fig. 7D, Figure S2 for the CyTOF datasets). Those changes were more significant than the case of the Influenza vaccination, possibly reflecting more intensive nature of the SARS-CoV-2 vaccine. Again, we examined and found that the strength and the timing of such responses depend on the original immune landscapes (see below for an exceptional case of H7). Notably, the initial induction of monocytes was generally higher with SARS-CoV-2, perhaps, consistent with the fact that inflammatory side effects of this vaccine, such as fever and inflammation, in this individual was stronger than the influenza vaccine. It should also be noted that, in this case, CD16^+^ non-classical monocytes were also induced, followed by the induction of CD14^+^ classical monocytes, indicating an enhanced immune response of this vaccination. In the second vaccination of H1 and H6 (Figs. 7E and 7F), the same response as mentioned above was observed.

We particularly attempted to inspect the changes of clonotypes after SARS-CoV-2 vaccination (sequencing stats shown in Table S7). The pre-vaccination TCR clonotypes were collectively removed. To particularly focus on the responses specific to the SARS-CoV-2 vaccine, all the VDJ sequences observed for influenza vaccination analysis were also subtracted (Figs. 7G and 7H). Consistent with influenza vaccination, the majority of the clonotypes were only detected at a single time point, reflecting the complexity of the TCR population. Nevertheless, a total of 20 and 62 clonotypes were identified from more than three-time points, in H1 and H6 datasets. Similar to influenza vaccination, characteristic sub-populations of the sporadic and persistent populations were also observed with SARS-CoV-2 vaccination (Figs. 7G and 7H, detailed in Figure S3).

To further characterize these sporadic and persistent populations, we examined the gene expression level of CD69 as a marker for activation of T cells. We found that the T cells of the persistent group showed higher CD69 levels, suggesting that those cells were in an active state (Figs. 7I and 7J). We further traced their time-lapse changes and identified several clonotypes that were induced at some time points. Most of those T cells were induced at the first vaccination and then further enhanced by the second vaccination (Figs. 7K and 7L). These TCRs may represent the cells that were specifically induced by this vaccination.

Interestingly, H7 showed an overall unique character. As described above, this individual originally had a higher percentage of NK cells. When we examined the changes in the immune cell profiles in H7 (Fig. 7D), the changes in the immune cell profiles were less relevant than H1 and H6. This observation may reflect the advanced age of this individual or a generally high level of NK cell-centered immune cell activity in its pristine state, possibly based on that individual’s medical history (Fig 1A, bottom, inset table). Although this individual eventually acquired a sufficient antibody level, the level obtained was, to some extent, lower than H1 and H6. Consistently, the changes of the CD69 levels were less significant, which in turn suggest that the vaccine responses depend on the original immune state of the individuals.

## DISCUSSION

In this study, we attempted to describe the diversity of the immune cell profiles in PBMC amongst healthy individuals. We revealed that the gene cellular components and gene expression profiles are diverse even in healthy individuals, possibly reflecting the personal history of previous immune responses. The unique point of this study is that we employed intensive scRNA-seq transcriptome and TCR VDJ sequencing analyses validated by single-cell mass spectrometry analysis. This approach could track a single clonotype across the sampling time points in association with its expressing T cell state. To the best of our knowledge, although there may be some previous studies which have analyzed immune cell profiles of healthy individuals, those studies used the Western population. In this study, we considered the Asian populations, which are supposed to show distinct immune responses to various pathogens, including SARS-CoV-2. By collecting and re-analyzing the previous data for the Western population and comparing it with the data of the present study, the immunological difference in health status will also be unveiled. Such insight is particularly of interest, considering that in the early stages of the COVID-19 pandemic, allegations were made that ethnicity may be responsible for the variability in susceptibility and morbidity worldwide(Barash et al. 2020; Sze et al. 2020; Bunyavanich et al. 2020; El-Khatib et al. 2020; Hou et al. 2020). Also. the difference in immune responses is also associated with the effectiveness of the vaccination. There are some papers describing the cause of such ethnic differences as the pre-existing discrepancies in health equity, such as access to healthcare and social determinants of health, and likely not genetics(Shelton et al. 2021; Lee et al. 2020). The ethnicity effects on immune response should also consider their medical recorts and the current enviromnents.

The obvious drawbacks of the present study include the general lack of in-depth biological validations. In particular, the results of the epitope identification were not validated for various pathogens. More generally, even after long discussions, the extent to which PBMC should represent the immune states of the individual remains debatable. Also, the small sample size and especially the short sampling period also limits the comparability of the results to previous in vitro laboratory studies. Particularly for the immune response to the vaccination, careful analyses are needed to elucidate what molecular events are occurring there in more detail. However, it is generally technically difficult to validate the unique events taking place in individuals overtime of their personal histories.

Nevertheless, it is significant that we could identify the individual heterogeneity of healthy immunity. The main aim of this study paper is to generate a base for such future studies to address the issues named above. In particular, the inter-individual heterogeneity was more pronounced than the intra-individual temporal variance. Therefore, future studies investigating immune system fluctuations in disease should account for the baseline diversity amongst healthy individuals, as demonstrated in this study.

No less important, we consider the present study results have indicated the importance of data collection for particular individuals. The immune cell profiles should be so diverse that in the event of a disease, the healthy state information should be directly subtracted, and the status of the immune cell analyzed. It is important to know the state of immune cells for infectious diseases and various types of other diseases, such as cancers. Recently, many anti-cancer drugs are designed to control proper or enhanced actions of the immune cells(June et al. 2018). Importantly, once the disease develops, the profile of the healthy states would be lost; thus, such information should be collected beforehand. This direction should be followed by the need for “personal immunological records.”. The records may include not only the data resource but also the banked biomaterial samples.

Personal health or medical histories, which differ depending on the immune responses experienced throughout their lives, should have collectively shaped their current immune landscapes. Such a landscape is the base to determine his or her unique immune condition in the daily life or to predict or control his or her response to various diseases.

Therefore, the “personal immune landscape” may become the data resource which should be prepared ideally for each individual. The present study should have paved the first step towards the new era of “personalized genomics” research and its social applications.

## METHODS

### Ethics approval and consent to participate

The human materials were collected and analyzed following the procedure approved by the ethical committee of the University of Tokyo as examination number: 20-351. All human subjects provided written informed consent.

### Library Preparation and Sequencing

PBMCs samples were collected from seven healthy donors (Table 1). Two participants, H1 and H2, had their PBMCs collected nine times over a month (Fig. 1A). A single sample was collected for the other participants. Except for the first sample from H1 and H2, samples were frozen and thawed before processing. Each sample was processed with the Chromium Next GEM Single Cell 5’ Library and Gel Bead Kit following the manufacturer’s user guide (10x Genomics, v1.1). After cDNA amplification, the B cell and T cell V(D)J were enriched using the human B cell and T cell enrichment kit before TCR and BCR library construction. The prepared 5’ GEX, TCR, and BCR libraries were then sequenced using the Illumina NovaSeq sequencer.

### Cell Type Annotation in sequencing dataset

The 5’ GEX dataset was initially processed with Cell Ranger (v.3.1.0 for the daily variation study, and version v5.1.0 for influenza and SARS-CoV-2 vaccination study), and underwent quality control and clustering by the R package Seurat (version 3.2 for daily variation study, and version 4.1 for influenza and SARS-CoV-2 vaccination study)(Stuart et al. 2019). Each cluster was primarily annotated on the differentially expressed gene set and the canonical markers retrieved from literature(Tian et al. 2019; Martos et al. 2020). T cells were identified based on *CD3D* and were determined as CD8^+^ or CD4^+^ depending on *CD8A* or *CD4* expression, respectively. CD8^+^ T cells were further classified into effector (*GZMK, GZMH, PRF1, CCL5*), memory (*CD29*) and naïve (*CCR7*). CD4^+^ T cells were similarly classified into naïve (*IL7R, CCR7*), memory (*IL7R, S100A4*), Treg (*FOXP3*). Other T cells included MAIT cells *(SLC4A10, TRAV1-2*) and γδT cells (*TRGV9, TRDV2*). B cells were identified if *MS4A1^+^* and were labeled naïve if *CD27-*. Plasma B cells were *MZB1^+^*/ *XBP1^+^.* Monocytes were grouped into either classical monocytes (*CD14, LYZ*), or non-classical monocytes (*FCGR3A, MS4A7*). DCs were typed as myeloid DCs (*FCER1A*, *CD1C*), and plasmacytoid DCs (*FCER1A, LILRA4*). We further identified NK cells (*GNLY, NKG7, CD56*) and platelets (*PPBP*). Additionally, SingleR (Aran et al. 2019) and Azimuth(Hao et al. 2021) was referenced to assist manual labeling where necessary. SingleR labels new cells from a test dataset based on similarity to a given reference dataset of samples with known labels, derived from either single-cell or bulk RNA-seq. This study used a publicly available bulk RNA-seq dataset of sorted immune cells as a reference for SingleR imputation(Monaco et al. 2019). Azimuth uses a precomputed supervised PCA (SPCA) transformation, a supervised version of principal component analysis to identify the best transcriptomic modules that delineate Weighted Nearest Neighbor-defined cell types.

### T cell Receptor and B cell Receptor Analysis

For scVDJ-seq of BCR and TCR, data were processed using Cell Ranger (v.3.1.0 for the daily variation study and version v5.1.0 for the influenza and SARS-CoV-2 vaccination study). For BCR analysis, data were analyzed by scRepertoire(Borcherding et al. 2020).

### Analysis using Mass Cytometry

We analyzed frozen PBMC samples using Helios Mass Cytometer (Fluidigm, sample list is shown in Table S17). We applied mass cytometry (CyTOF) using the Maxpar® DirectTM Immune Profiling AssayTM to characterize PBMCs. Samples are processed following vendors’ guide (Quick Reference guide JPN_PN 400288 B1_001). Briefly, counted PBMCs are washed with cell staining buffer and processed with FcX. Cells are stained with 30 antibodies shown in Key resources table. After cell staining, cells are fixed with 1.6% formaldehyde. Stained cells are analyzed with Helios. Datasets were analyzed with Maxpar Pathsetter.

### Antibody Titration

In the influenza vaccination study, we measured the antibody titer for type A-H1, type A-H3, type B-Yamagata, and type B-Victoria flu using HI method. In the SARS-CoV-2 vaccination study, we measured the antibody titer anti-SARS-CoV-2 S IgG and anti-SARS-CoV-2 S IgM.

### Intracellular cytokine staining assay using Flow Cytometry

In the intracellular cytokine staining assay, we stimulated PBMC of H1, H2 and H3 with CMV (pp65 and IE1), EBV (EBNA1, LMP1, and BZLF1) and incubated for six hours in 37C. For staining, we used FITC (CD4), PE (CD107a), PerCP (CD8a), PE-Cy7 (IL2), APC (TNFa), APC-Cy7 (IFNg), Pacific Blue (CD3) and LIVE/DEAD Aqua-Amcyan Antigen for staining cells. We employed BD FACSCanto (BD) and the sort logic was set by gating lymphocytes by forward scatter and side scatter and then gating on CD3^+^ CD4^+^ cells and CD3^+^ CD8^+^ cells. The dataset was analyzed by FlowJo sofetware.

## DATA ACCESS

The raw data has been deposited to National Bioscience Database Center as study number: JGAS000321. The present study did not develop any new software. All code used in the present study can be available upon request to Lead Contact, Yutaka Suzuki.

## COMPETEING INTEREST STATEMENT

The authors decleare that no competing interests exits.

### Author’s Contributions

Y.K. performed scRNA-seq experiment, visualizeds scRNA-seq and CyTOF results and drafted the manuscript. P.R. performed CyTOF experiment. N.Y. performed the analysis of scRNA-seq. L.R., S.N., and M.O. were involved in blood sample collection and contributed to drafting manuscripts. K.K., A.S., M.Seki, M.Sakata, Y.I., A.K.T., K.H.N., and T.M. contributed to drafting manuscripts. Y.S. supervised the project. All authors reviewed, approved, and accepted the manuscript.

## Acknowledgments

We thank all the anonymous donors who participated in this study. We appreciate Shintaro Yanagimoto for supporting immunological studies. We thank K. Imamura, K. Abe, Y. Ishikawa, M. Konbu, E. Kobayashi, E. Ishikawa, and S. Minamiguchi, Y. Kuze for their technical assistance.

## Additional files

Supplemental information includes three figures (separate file) and eleven tables (sepatarate file).

**Table S1 (separate file) Datasets used in the current study**

This table provides the list of all samples and methods used for analyses in this study.

**Table S2 (separate file) Sequence Statistics of scRNA-seq used in the daily and individual variance**

This table provides the sequence statistics of scRNA-seq 5’GEX for daily variance, individual variance, influenza vaccination and SARS-CoV-2 vaccination study.

**Table S3 (separate file) Percentage of simple cell type analyzed by scRNA-seq**

This table provides the percentage of the simple cell type for each sample.

**Table S4 (separate file) Percentage of detailed cell type analyzed by scRNA-seq**

This table provides the percentage of the simple cell type for each sample.

**Table S5 (separate file) List of genes with low-correlation value in fresh and frozen comparison**

This table shows the correlation of gene expression between fresh and frozen H1 Day 0 samples.

**Table S6 (separate file) List of correlation value in scRNA-seq and CyTOF comparison**

This table shows the correlation of scRNA-seq and CyTOF in the indicated cell types.

**Table S7 (separate file) Sequence statistics scVDJ-seq used in the current study**

This table provides the sequence statistics of the scRNA-seq TCR and BCR V(D)J sequencing of daily variance, individual variance, influenza vaccination and SARS-CoV-2 vaccination study.

**Table S8 (separate file) Frequency of top used Clonotypes in H1 and H2 TCR This table provides the statistics of the CDR3 sequence and the frequency of the daily variation of H1 and H2 study.**

**Table S9 (separate file) Statistics of the clonotypes and T cell recognizing CMV and EBV**

This table provides the information about the clonotypes and T cell recognizing CMV **(A)** and EBV **(B)**.

**Table S10. (separate file) Detected antibody level during influenza vaccination**

(A) This table shows the antibody level of the anti-influenza virus, type A-H1, type A-H3, type B-Yamagata, and type B-Victoria, during the influenza vaccination in H1 and H2. (B) This table shows the level of IgG and IgM antibody anti-SARS-CoV-2 virus in the S region during the SARS-CoV-2 vaccination in H1, H6 and H7.

**Table S11. (separate file) Markers used in the CyTOF analysis**

This table provides the markers used in the CyTOF study. We used a commercial antibody set.

## References

1. Adams NM, Grassmann S, Sun JC. 2020. Clonal expansion of innate and adaptive lymphocytes. Nat Rev Immunol 20: 694–707. http://www.nature.com/articles/s41577-020-0307-4.

2. Aran D, Looney AP, Liu L, Wu E, Fong V, Hsu A, Chak S, Naikawadi RP, Wolters PJ, Abate AR, et al. 2019. Reference-based analysis of lung single-cell sequencing reveals a transitional profibrotic macrophage. Nat Immunol 20: 163–172. http://www.ncbi.nlm.nih.gov/pubmed/30643263.

3. Barash A, Machluf Y, Ariel I, Dekel Y. 2020. The Pursuit of COVID-19 Biomarkers: Putting the Spotlight on ACE2 and TMPRSS2 Regulatory Sequences. Front Med 7: 582793. http://www.ncbi.nlm.nih.gov/pubmed/33195331.

4. Borcherding N, Bormann NL, Kraus G. 2020. scRepertoire: An R-based toolkit for single-cell immune receptor analysis. F1000Research 9: 47. http://www.ncbi.nlm.nih.gov/pubmed/32789006.

5. Bunyavanich S, Grant C, Vicencio A. 2020. Racial/Ethnic Variation in Nasal Gene Expression of Transmembrane Serine Protease 2 (TMPRSS2). JAMA 324: 1567–1568. http://www.ncbi.nlm.nih.gov/pubmed/32910146.

6. De Kouchkovsky I, Abdul-Hay M. 2016. “Acute myeloid leukemia: a comprehensive review and 2016 update”. Blood Cancer J 6: e441. http://www.ncbi.nlm.nih.gov/pubmed/27367478.

7. El-Khatib Z, Jacobs GB, Ikomey GM, Neogi U. 2020. The disproportionate effect of COVID-19 mortality on ethnic minorities: Genetics or health inequalities? EClinicalMedicine 23: 100430. http://www.ncbi.nlm.nih.gov/pubmed/32572393.

8. Ewer KJ, Barrett JR, Belij-Rammerstorfer S, Sharpe H, Makinson R, Morter R, Flaxman A, Wright D, Bellamy D, Bittaye M, et al. 2021. T cell and antibody responses induced by a single dose of ChAdOx1 nCoV-19 (AZD1222) vaccine in a phase 1/2 clinical trial. Nat Med 27: 270–278. http://www.ncbi.nlm.nih.gov/pubmed/33335323.

9. Gielis S, Moris P, Bittremieux W, De Neuter N, Ogunjimi B, Laukens K, Meysman P. 2019. Detection of Enriched T Cell Epitope Specificity in Full T Cell Receptor Sequence Repertoires. Front Immunol 10: 2820. http://www.ncbi.nlm.nih.gov/pubmed/31849987.

10. Golubovskaya V, Wu L. 2016. Different Subsets of T Cells, Memory, Effector Functions, and CAR-T Immunotherapy. Cancers (Basel) 8. http://www.ncbi.nlm.nih.gov/pubmed/26999211.

11. Hao Y, Hao S, Andersen-Nissen E, Mauck WM, Zheng S, Butler A, Lee MJ, Wilk AJ, Darby C, Zager M, et al. 2021. Integrated analysis of multimodal single-cell data. Cell 184: 3573–3587.e29. http://www.ncbi.nlm.nih.gov/pubmed/34062119.

12. Hayes MP, Berrebi GA, Henkart PA. 1989. Induction of target cell DNA release by the cytotoxic T lymphocyte granule protease granzyme A. J Exp Med 170: 933–46. http://www.ncbi.nlm.nih.gov/pubmed/2788710.

13. Hou Y, Zhao J, Martin W, Kallianpur A, Chung MK, Jehi L, Sharifi N, Erzurum S, Eng C, Cheng F. 2020. New insights into genetic susceptibility of COVID-19: an ACE2 and TMPRSS2 polymorphism analysis. BMC Med 18: 216. http://www.ncbi.nlm.nih.gov/pubmed/32664879.

14. June CH, O’Connor RS, Kawalekar OU, Ghassemi S, Milone MC. 2018. CAR T cell immunotherapy for human cancer. Science (80-) 359: 1361–1365. https://www.sciencemag.org/lookup/doi/10.1126/science.aar6711.

15. Lee I-H, Lee J-W, Kong SW. 2020. A survey of genetic variants in SARS-CoV-2 interacting domains of ACE2, TMPRSS2 and TLR3/7/8 across populations. Infect Genet Evol 85: 104507. http://www.ncbi.nlm.nih.gov/pubmed/32858233.

16. Martos SN, Campbell MR, Lozoya OA, Wang X, Bennett BD, Thompson IJB, Wan M, Pittman GS, Bell DA. 2020. Single-cell analyses identify dysfunctional CD16+ CD8 T cells in smokers. Cell reports Med 1. http://www.ncbi.nlm.nih.gov/pubmed/33163982.

17. Minervina A, Pogorelyy M, Mamedov I. 2019. T-cell receptor and B-cell receptor repertoire profiling in adaptive immunity. Transpl Int 32: 1111–1123. http://www.ncbi.nlm.nih.gov/pubmed/31250479.

18. Monaco G, Lee B, Xu W, Mustafah S, Hwang YY, Carré C, Burdin N, Visan L, Ceccarelli M, Poidinger M, et al. 2019. RNA-Seq Signatures Normalized by mRNA Abundance Allow Absolute Deconvolution of Human Immune Cell Types. Cell Rep 26: 1627–1640.e7. http://www.ncbi.nlm.nih.gov/pubmed/30726743.

19. Nicholson LB. 2016. The immune system. Essays Biochem 60: 275–301. http://www.ncbi.nlm.nih.gov/pubmed/27784777.

20. Olson TL, Cheon H, Xing JC, Olson KC, Paila U, Hamele CE, Neelamraju Y, Shemo BC, Schmachtenberg MW, Sundararaman SK, et al. 2021. Frequent Somatic TET2 Mutations in Chronic NK-LGL Leukemia with Distinct Patterns of Cytopenias. Blood. https://ashpublications.org/blood/article/doi/10.1182/blood.2020005831/475646/Frequent-Somatic-TET2-Mutations-in-Chronic-NK-LGL.

21. Osińska I, Popko K, Demkow U. 2014. Perforin: an important player in immune response. Cent J Immunol 39: 109–15. http://www.ncbi.nlm.nih.gov/pubmed/26155110.

22. Shelton JF, Shastri AJ, Ye C, Weldon CH, Filshtein-Sonmez T, Coker D, Symons A, Esparza-Gordillo J, 23andMe COVID-19 Team, Aslibekyan S, et al. 2021. Trans-ancestry analysis reveals genetic and nongenetic associations with COVID-19 susceptibility and severity. Nat Genet 53: 801–808. http://www.ncbi.nlm.nih.gov/pubmed/33888907.

23. Shi L, Kraut RP, Aebersold R, Greenberg AH. 1992. A natural killer cell granule protein that induces DNA fragmentation and apoptosis. J Exp Med 175: 553–66. http://www.ncbi.nlm.nih.gov/pubmed/1732416.

24. Shugay M, Bagaev D V, Zvyagin I V, Vroomans RM, Crawford JC, Dolton G, Komech EA, Sycheva AL, Koneva AE, Egorov ES, et al. 2018. VDJdb: a curated database of T-cell receptor sequences with known antigen specificity. Nucleic Acids Res 46: D419–D427. http://www.ncbi.nlm.nih.gov/pubmed/28977646.

25. Spitzer MH, Nolan GP. 2016. Mass Cytometry: Single Cells, Many Features. Cell 165: 780–91. http://www.ncbi.nlm.nih.gov/pubmed/27153492.

26. Stuart T, Butler A, Hoffman P, Hafemeister C, Papalexi E, Mauck WM, Hao Y, Stoeckius M, Smibert P, Satija R. 2019. Comprehensive Integration of Single-Cell Data. Cell 177: 1888–1902.e21. https://linkinghub.elsevier.com/retrieve/pii/S0092867419305598.

27. Stubbington MJT, Rozenblatt-Rosen O, Regev A, Teichmann SA. 2017. Single-cell transcriptomics to explore the immune system in health and disease. Science 358: 58–63. http://www.ncbi.nlm.nih.gov/pubmed/28983043.

28. Sze S, Pan D, Nevill CR, Gray LJ, Martin CA, Nazareth J, Minhas JS, Divall P, Khunti K, Abrams KR, et al. 2020. Ethnicity and clinical outcomes in COVID-19: A systematic review and meta-analysis. EClinicalMedicine 29: 100630. http://www.ncbi.nlm.nih.gov/pubmed/33200120.

29. Tian Y, Grifoni A, Sette A, Weiskopf D. 2019. Human T Cell Response to Dengue Virus Infection. Front Immunol 10: 2125. http://www.ncbi.nlm.nih.gov/pubmed/31552052.

30. Verhoeckx K, Cotter P, López-Expósito I, Kleiveland C, Lea T, Mackie A, Requena T, Swiatecka D, Wichers H, eds. 2015. The Impact of Food Bioactives on Health. Springer International Publishing, Cham http://link.springer.com/10.1007/978-3-319-16104-4.

31. Vivier E, Tomasello E, Baratin M, Walzer T, Ugolini S. 2008. Functions of natural killer cells. Nat Immunol 9: 503–510. http://www.nature.com/articles/ni1582.

32. Weese D, Holtgrewe M, Reinert K. 2012. RazerS 3: faster, fully sensitive read mapping. Bioinformatics 28: 2592–9. http://www.ncbi.nlm.nih.gov/pubmed/22923295.

33. Zhang J-Y, Wang X-M, Xing X, Xu Z, Zhang C, Song J-W, Fan X, Xia P, Fu J-L, Wang S-Y, et al. 2020. Single-cell landscape of immunological responses in patients with COVID-19. Nat Immunol 21: 1107–1118. http://www.ncbi.nlm.nih.gov/pubmed/32788748.

